# Active maintenance of meiosis-specific chromosome structures in *C. elegans* by the deubiquitinase DUO-1

**DOI:** 10.1101/2025.09.11.675685

**Authors:** Liesl G. Strand, Charlotte P. Choi, Savannah McCoy, Emmanuel T. Nsamba, Nicola Silva, Anne M. Villeneuve

**Affiliations:** Department of Developmental Biology, Stanford University School of Medicine, Stanford, California, USA; Department of Biology, Masaryk University, Faculty of Medicine, Brno, Czech Republic; Department of Genetics, Stanford University School of Medicine, Stanford, California, USA

## Abstract

Meiotic prophase is characterized by a dynamic program in which germ cells undergo a complex series of associations and dissociations of protein complexes that drive assembly, remodeling, and disassembly of meiosis-specific chromosome structures and dramatic changes in chromosome compaction. Failure to properly coordinate these processes can result in improper chromosome segregation, producing aneuploid gametes and inviable zygotes. Here, we investigate the roles of *C. elegans* DUO-1, an ortholog of mammalian ubiquitin-specific proteases USP26 and USP29, in mediating these dynamic chromosomal events during meiotic prophase. Cytological analyses of *duo-1* null mutants indicate that loss of DUO-1 function leads to impaired assembly of synaptonemal complexes (SCs), loss of integrity of meiotic chromosome axes, ineffective homolog pairing, premature separation of sister chromatids, and late-prophase chromosome decompaction. Further, SC instability in *duo-1* mutants correlates with depletion of REC-8 cohesin complexes and is accompanied by massive accumulation of early DSB repair intermediates. By using a dual-AID-tagged allele to deplete DUO-1 during meiotic development, we demonstrate that DUO-1 is continually required throughout meiotic prophase progression, to promote proper axis/SC assembly in early prophase, to maintain axis/SC stability during the late pachytene stage, and to promote/maintain chromosome compaction at the end of meiotic prophase. Together, our data reveal that meiotic chromosome structure and meiosis-specific chromosome architecture require active maintenance throughout meiotic prophase, and that this maintenance is necessary for successful meiosis.

## Introduction

During meiotic prophase, germ cells undergo multiple major state transitions alongside dramatic remodeling of chromosome architecture and nuclear organization *en route* to creating the haploid gametes that are essential for sexual reproduction. Upon meiotic prophase entry, chromosomes acquire meiosis-specific architecture in which replicated sister chromatids become organized into a series of chromatin loops associated at their bases with a highly-ordered meiosis-specific structure known as the chromosome axis (reviewed in ref. 1). Meiosis-specific cohesion complexes are key architectural components of the axis, with some complexes mediating cohesion between sister chromatids, while others organize the chromatin into axis-associated loops. This cohesion-based organization in consolidated into coherent axial structures through the incorporation of additional meiosis-specific proteins. Notable among these are members of the conserved, yet rapidly evolving, meiosis-specific HORMA domain protein family, which have the capacity to interact with each other, with additional axis components and/or with chromatin in a manner that yields long extended multimeric assemblies.

As homologous chromosome pairing progresses, these axial structures become the lateral elements of a tripartite structure known as the synaptonemal complex (SC) (2) which assembles between co-aligned homologs. Association between axial proteins and components of the central region of the SC creates a highly organized, yet dynamic, interface between homologous chromosomes that serves as an environment for organizing the DNA events of meiotic recombination (1, 3). The meiotic axis is implicated in promoting and regulating formation of double-strand DNA breaks (DSBs), which serve as the initiating events of meiotic recombination. SC central region proteins are further implicated in promoting conversion of a subset of these DSBs into crossovers (COs) and/or in regulating the number of COs and their distribution along chromosomes.

Following completion of CO formation, chromosomes undergo another major structural remodeling organized around CO sites, involving SC disassembly and reorganization of chromosome axes. This remodeling is typically followed by further condensation and compaction to yield highly compact bivalents in which homologous chromosomes are physically connected yet oriented away from each other in a manner that enables attachment to and segregation toward opposite poles of the meiosis I spindle (4).

The nematode *C. elegans* provides an outstanding experimental system for investigating the dynamic organization of chromosome structure during meiotic prophase, as the events under study are highly accessible to cytological analyses in the context of a transparent gonad in which germ cells are arranged in a developmental time course of progression though meiotic prophase (5). Moreover, the known inventory of structural components of meiotic chromosome axes and the SC central region is extensive. *C. elegans* contains two distinct meiosis-specific cohesion complexes that are interspersed along the meiotic chromosome axes, with REC-8 cohesin complexes functioning primarily to mediate sister chromatid cohesion and COH-3/4 cohesin complexes functioning primarily in loop formation (6–10). Four different meiotic HORMAD proteins (HTP-3, HTP-1/2 and HIM-3) assemble in a hierarchical manner, together conferring multiple structural and functional properties of axes, *e.g.* in assembling SCs, in promoting DSB formation, in surveillance of the status of chromosomal events as part of a crossover assurance checkpoint, and in creating differentiated axis sub-domains to enable two-step loss of sister chromatid cohesion during the meiotic division (9, 11–18). A comprehensive set of SC central region proteins has also been identified, comprising six coiled-coil-containing proteins (SYP-1, SYP-2, SYP-3, SYP-4 and SYP-5/6) that are interdependent for chromosomal loading and stability (19–24) and two paralogous proteins (SKR-1 and SKR-2) that are orthologs of SCF ubiquitin ligase adaptor Skp1 but play a “moonlighting” role as integral structural components of the SC (25). Finally, mutagenesis-based and RNAi-based genetic screening approaches in *C. elegans* have been highly effective for discovering numerous key components of the meiotic machinery (5, 26, 27). These approaches have not only identified multiple essential meiotic recombination proteins and many of the structural components discussed above, they have also identified important factors that coordinate and regulate meiotic events to ensure orderly progression and enforce quality control to achieve a successful outcome of the meiotic divisions.

In the current work, we report on our discovery of *C. elegans* DUO-1, an ortholog of mammalian ubiquitin-specific proteases USP26 and USP29, as a factor essential for the success of multiple distinct aspects of meiotic prophase chromosome dynamics. By analyzing the consequences of both systemic loss and short-term depletion of DUO, we demonstrate that DUO-1 is required to promote proper assembly of SC between homologous chromosome pairs, to prevent accumulation of early recombination intermediates, to maintain cohesion between sister chromatids and structural integrity of the SC, and to promote and maintain compaction of meiotic bivalents during late prophase. Together, our findings reveal that meiosis-specific chromosome structures and chromosome architecture require active maintenance throughout meiotic prophase to achieve a successful outcome of sexual reproduction.

## Results

### Identification of DUO-1 as a component of the *C. elegans* meiotic machinery

We were prompted to investigate the roles of DUO-1 in meiosis following isolation of several independent *duo-1* mutant alleles in recent genetic screens for mutants defective in meiotic chromosome segregation (Fig 1A). In these screens, adapted from the “Green eggs & Him approach of Kelly et al. (2000), hermaphrodites homozygous for *duo-1* mutations were identified based on production of embryos expressing a GFP reporter indicative of X-chromosome mis-segregation (26). After using CRISPR-based allele reconstruction to confirm that the identified *duo-1* point mutations were causative for meiotic defects, we generated a STOP-IN (28) null allele, *duo-1(me186),* to characterize the meiotic roles of DUO-1 in more detail.

**Figure 1.**
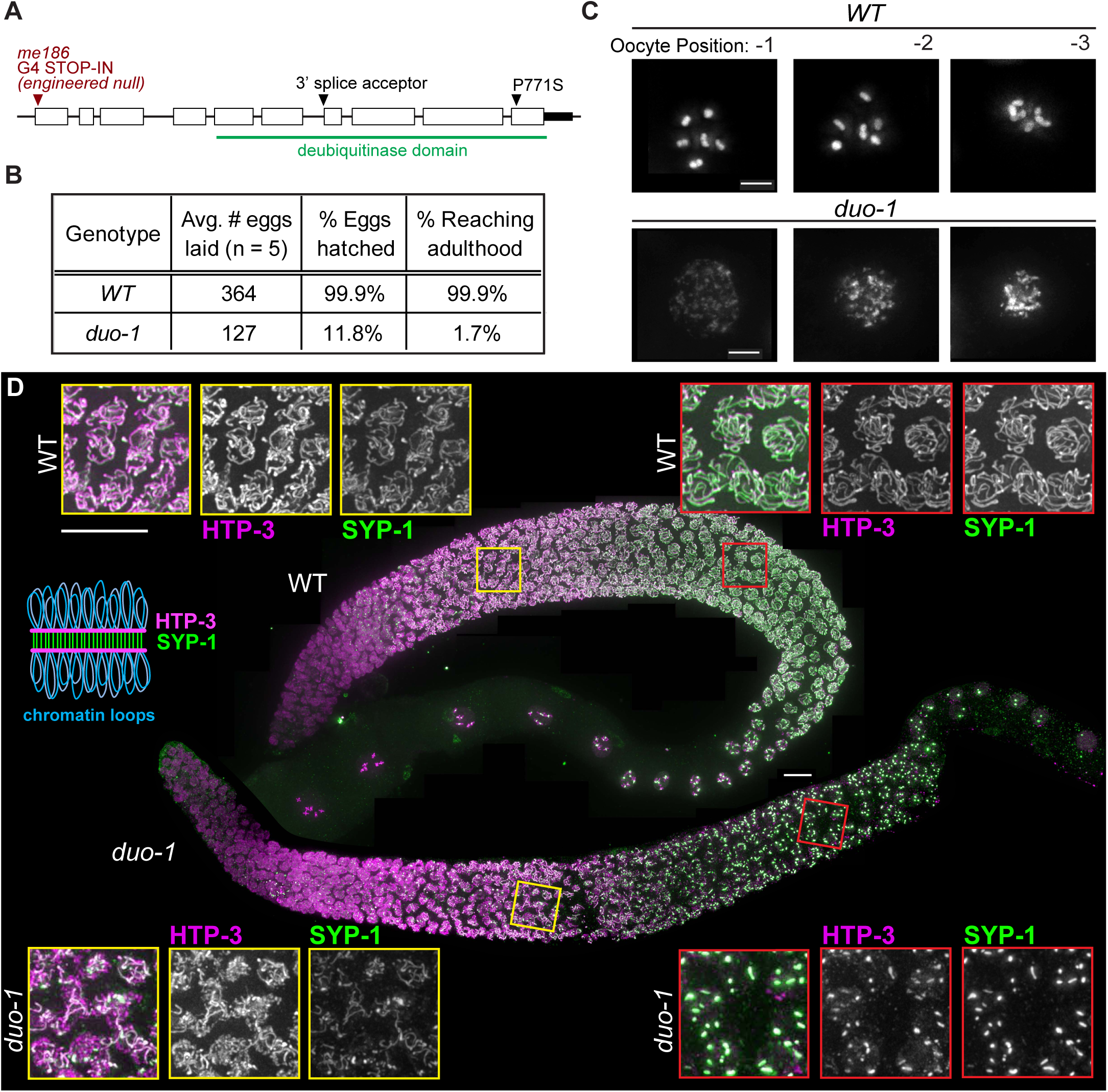
DUO-1 is required for proper meiosis. *A) duo-1* gene structure, with the positions and nature of mutations described in this work. The *me186* CRISPR-engineered null allele is referred to as *duo-1* throughout the text. *B)* Quantification of decreased brood size, hatch rate, and fraction of progeny reaching adulthood in *duo-1* mutants. *C)* DAPI-stained proximal oocytes in WT and *duo-1* animals, showing late diakinesis decompaction phenotype in *duo-1* mutants. Numbers indicate location of oocyte relative to spermatheca. Scale bar = 5µm. *D)* Max-projection immunofluorescence images of whole-mount WT and *duo-1* gonads stained for axis component HTP-3 and synaptonemal complex (SC) central region component SYP-1, as indicated in the schematic diagram. Insets show greyscale images of HTP-3 and SYP-1 in the early (left) and late (right) pachytene regions of WT and *duo-1* mutant germlines, illustrating partial assembly of SC components along chromosomes in early pachytene and SC-proteins concentrated into large aggregates by late pachytene in the *duo-1* mutant. Scale bars = 10µm.

Notably, *duo-1* mutants exhibit highly decompacted chromatin in oocytes at diakinesis, the last stage of meiotic prophase, when chromosomes are normally condensed into compact bivalents in preparation for the meiotic divisions (Fig 1C). This unusual phenotype, in which highly decondensed chromatin is distributed throughout the nucleus and chromosomes are not resolvable as separate entities, differs from the vast majority of *C. elegans* meiotic mutants, in which compact but unconnected chromosomes (univalents) are observed in diakinesis nuclei (29–32).

Most embryos produced by *duo-1* mutant hermaphrodites fail to hatch, consistent with expectations for mutants with defective meiosis (Fig 1B). However, the fraction of embryos that hatch (12%) is surprisingly high given the aberrant state of chromosomes in diakinesis oocytes (though most hatched larvae develop slowly and/or do not reach adulthood) and is higher than typically observed for meiotic mutants with compact univalents at diakinesis (3-5%; e.g. refs. 22, 30). To investigate the possibility that this higher hatching frequency might reflect successful spermatocyte meiosis, we conducted cytological analyses of gonads from *duo-1* mutant males; however, these analyses revealed that spermatocyte meiosis is also impaired, as evidenced by unusual morphology and lack of uniformity of DAPI signals in post-meiotic spermatids (Fig S1). We conclude that impairment of meiotic chromosome segregation in *duo-1* mutants likely yields different patterns of aneuploidy than occurs in most other meiotic mutants, where chromosomes are properly condensed but unconnected.

### DUO-1 is required for assembly/stability of synaptonemal complexes

In light of the aberrant organization of chromosomes in diakinesis oocytes in *duo-1* mutants, we evaluated the effects of loss of *duo-1* function on the state of meiosis-specific chromosome structures throughout meiotic prophase. We used immunofluorescence microscopy to visualize components of the synaptonemal complex (SC), which forms at the interface between homologous chromosomes during meiotic prophase and is a prerequisite for normal bivalents at diakineses. Figure 1D shows immunostaining for axial element protein HTP-3 and SC central region protein SYP-1. In wild-type hermaphrodite germ lines, HTP-3 becomes visible as axial structures upon meiotic prophase entry, and SYP-1 colocalizes with HTP-3 as the SC assembles at the interface between aligned pairs of homologous chromosomes (11, 22, 33–35). SYP-1 and HTP-3 remain colocalized along the lengths of SCs throughout the pachytene stage, with SYP-1 becoming enriched toward one end of each SC just prior to SC disassembly.

In *duo-1* mutant germ lines, HTP-3 is also detected in axis-like structures in early prophase nuclei, and SYP-1 colocalizes with a subset of the HTP-3 signal. However, nuclei fail to complete SC assembly, and SYP-1 does not become fully localized along HTP-3 tracks. Incomplete SC assembly in the *duo-1* mutants is accompanied by an extension of the polarized chromatin distribution characteristic of transition zone nuclei, indicating a failure to satisfy checkpoint requirements for meiotic progression in a timely manner (Fig S2). Moreover, as prophase progresses, these assembled structures break down, and HTP-3 and SYP-1 instead become colocalized into large aggregates reminiscent of the polycomplexes that form in mutants lacking essential components of chromosome axes (11, 36–38). These aggregates persist through the region of the germ line corresponding to the diplotene stage in wild-type meiosis.

This pattern of partial SC assembly followed by SC breakdown is also observed in *duo-1* germ lines stained for axial element component HIM-3 (39) (Fig S2). Further, the RING finger protein ZHP-3, which is normally recruited to the SC central region by the SYP proteins (31, 40), likewise colocalizes with HIM-3 tracks in early prophase and later in aggregates. *duo-1* mutant male germlines display a very similar SC phenotype (Fig S1). We conclude that while SC assembly can be initiated in *duo-1* mutants, DUO-1 is required for full assembly and stability of the nascent SC.

### DUO-1 is required for normal pairing of homologous chromosomes and for maintenance of sister chromatid cohesion

As SC assembly normally stabilizes associations between homologous chromosome pairs, we evaluated homolog pairing in *duo-1* mutant germ lines. Figure 2A shows a time course analysis in which pairing was assessed by *fluorescence in situ hybridization* (FISH) using a set of probes targeting a 1Mb region on chromosome II (Fig 2; Methods). In wild-type germ lines, two separate FISH signals were detected in most nuclei prior to meiotic entry (reflecting unpaired homologs), whereas soon after the onset of meiotic prophase and continuing through to the end of the pachytene stage most nuclei had a single FISH signal, reflective of paired homologs. Premeiotic nuclei in the *duo-1* mutant likewise exhibited two FISH signals. Following meiotic entry, however, most *duo-1* mutant nuclei continued to exhibit two FISH signals and the incidence of nuclei with a single FISH signal did not increase, indicating a severe impairment of homolog pairing. Moreover, in the region of the *duo-1* germ line corresponding to very late pachytene/transition to diplotene in the wild type, three or four FISH signals were detected in the majority of nuclei. Presence of nuclei with greater than two FISH signals is indicative of a premature breakdown of cohesion between sister chromatids in the *duo-1* mutant.

**Figure 2.**
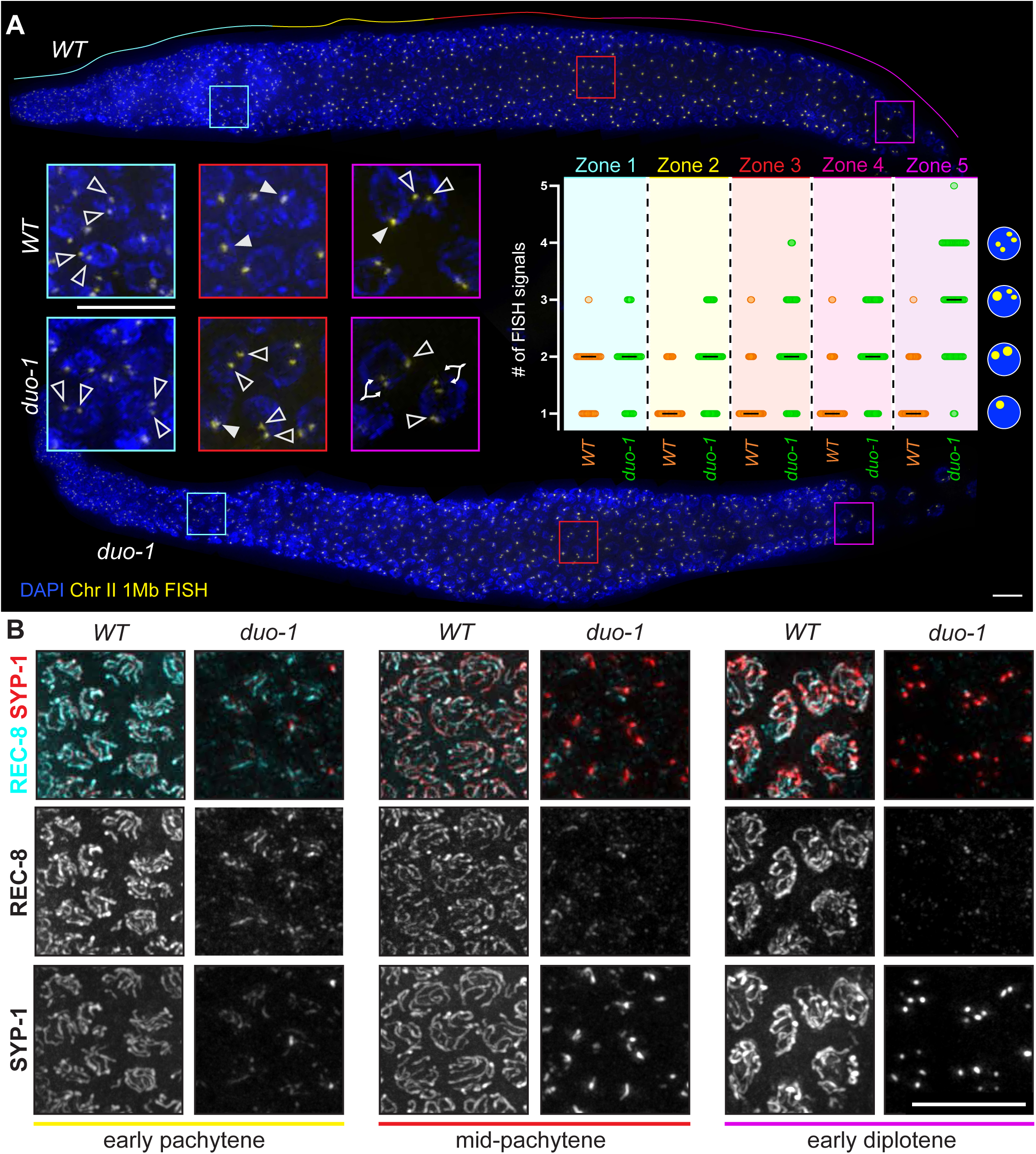
DUO-1 is required for normal homolog pairing and for maintenance of sister chromatid cohesion. *A)* Fluorescence *in situ* hybridization (FISH) images of whole-mount gonads showing defective pairing and chromatid cohesion in *duo-1* mutants for a locus on chromosome II. Insets show magnified nuclei from the pre-meiotic (cyan), mid-pachytene (red), and late pachytene/early diplotene (magenta) regions. Open arrowheads indicate FISH signals in nuclei with unpaired homologous chromosomes; filled arrowheads point to a single FISH signal within a nucleus, indicative of paired homologs; branched arrowheads show two close-set signals that represent separated sister chromatids in nuclei with 3-4 foci, as indicated in schematic at right of graph. Graph shows quantification of FISH signals in nuclei in consecutive zones (indicated by color) along the gonad axis (see Methods for full description of zone delineation). Each circle represents a nucleus; black lines indicate median number of FISH signals observed. For WT, Mann-Whitney tests indicate a significant difference between Zone 1 and Zone 2 (p <0.0001) but no significant differences between other adjacent zones within WT, indicating that chromosome II homologs become paired and stay paired. For *duo-1*, Zone 4 and Zone 5 were significantly different from each other (p <0.0001) but no other significant differences were detected, indicating that *duo-1* mutants fail to pair chromosome II and then prematurely lose sister chromatid cohesion in late pachytene. Comparing the same zones between *duo-1* and WT yielded the following P values: 0.2028 (ns, Zone 1) and <0.0001 (Zone 2, Zone 3, Zone 4, and Zone 5). Scale bars = 10µm. *B)* Insets from max-projected immunofluorescence images of whole-mount gonads, showing depletion of cohesin complex component REC-8 from chromosomes in the *duo-1* mutant. While SYP-1 and REC-8 colocalize on partially assembled axes in early prophase in *duo-1* mutants, REC-8 is lost from chromosomes as SCs break down and is only detected in a subset of SC-aggregates in the mid-pachytene and early diplotene regions. Scale bar = 10µm.

Loss of *duo-*1 function does not result in a complete loss of homolog pairing, however, as successful pairing at the *X* chromosome pairing center is detected in the *duo-1* mutant by immunolocalization of pairing center binding protein HIM-8 (41) (Fig S3). Differences between X-chromosomes and autosomes in genetic requirements for pairing and synapsis are well known in *C. elegans* and are associated with differences in timing of replication and synapsis, chromatin state, and chromosome compaction (14, 15, 41–43). We note that the more compact nature of the X chromosomes relative to autosomes is still evident in the *duo-1* mutant through the late pachytene region of the germ line, based on the appearance of DAPI-stained chromatin adjacent to HIM-8 signals (Fig S3).

The premature loss of sister-chromatid cohesion detected by FISH analysis in *duo-1* mutants suggests a failure of the cohesin complexes that normally hold sisters together throughout prophase I. We confirmed this expectation through immunolocalization of REC-8, the kleisin subunit of the meiosis-specific cohesin complexes that are responsible for mediating sister-chromatid cohesion during *C. elegans* meiosis (7, 8, 10) (Fig 2B). As a component of meiotic chromosome axes, REC-8 exhibits axis/SC localization throughout most of meiotic prophase during wild-type meiosis. In *duo-1* mutants, REC-8 is also initially detected in axis stretches, colocalizing with SYP-1 in regions of partial synapsis, but REC-8 signal becomes diminished as SCs disassemble during prophase progression and SC proteins become aggregated in polycomplexes. These data indicate that REC-8 cohesin complexes are no longer distributed along chromosomes in a manner that could maintain sister cohesion.

Assessment of polar body number in post-meiotic early embryos provides further evidence for premature loss of REC-8-mediated cohesion in *duo-1* mutants (Fig S4). Whereas two extruded polar bodies (reflecting successful completion of the two oocyte meiotic divisions) are detected in most wild-type embryos, most *duo-1* mutant embryos have only a single polar body. This single polar body phenotype is a hallmark of mutants that lack or exhibit premature loss of REC-8 (44), reflecting equational segregation of sister chromatids occurring at meiosis I coupled with failure of polar body extrusion at meiosis II.

### DUO-1 is required to prevent accumulation of early DSB repair intermediates

Meiotic chromosome axes and SCs are intimately associated with formation and repair of double-strand DNA breaks (DSBs), which serve as the initiating events for meiotic recombination (45). In light of the disruption of axis/SC organization in *duo-1* mutant nuclei, we used immunofluorescence to assess the appearance and/or disappearance of foci corresponding to DSB repair proteins localized at recombination sites.

As depicted in Fig 3A, we found that *duo-1* mutants exhibit a major alteration in the localization dynamics of recombinase RAD-51, a DNA strand exchange protein that associates with early recombination intermediates formed by resection of DSB ends (20, 46). RAD-51 foci begin to appear with similar timing in wild-type and *duo-1* mutant germ lines, rising in abundance during early pachytene. However, while RAD-51 foci decline in number during the mid-pachytene stage and disappear by the late pachytene stage during wild-type meiosis, RAD-51 signals instead intensify and foci continue to increase in numbers in the mid-to late pachytene region of *duo-1* mutant germ lines and remain highly abundant into later prophase, reflecting hyperaccumulation of RAD-51 (Fig 3B-C). Importantly, this marked accumulation of RAD-51 is dependent on the DSB-generating enzyme SPO-11 (29), as RAD-51 foci are abolished in *spo-11; duo-1* germ lines (Fig S5), indicating that RAD-51 accumulation represents persistence and/or accumulation of RAD-51 at meiotic DSB repair intermediates.

**Figure 3.**
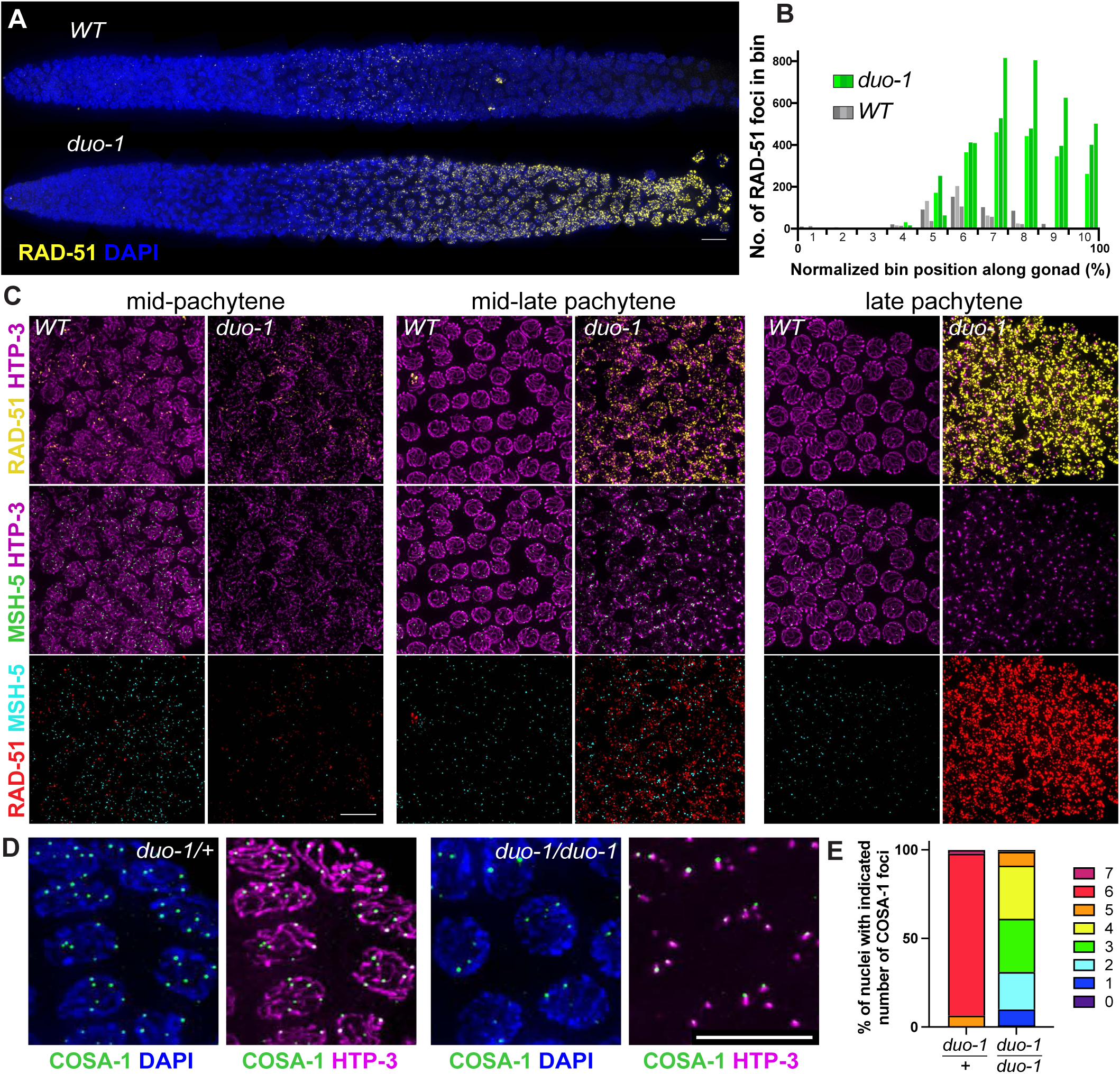
DUO-1 is required to prevent accumulation of early DSB repair intermediates. *A)* Straightened images of whole-mount gonads immunostained for DNA strand exchange protein RAD-51, which localizes to early DSBR sites, showing hyperaccumulation of RAD-51 in the *duo-1* mutant. Scale bar = 10µm. *B)* Quantification of RAD-51 foci in ten bins evenly distributed across the length of the gonad (see Methods). Three gonads (represented by different shades of grey or green) were quantified for each genotype. *C)* Nuclear spreads costained for RAD-51, HTP-3, and the meiotic recombination factor MSH-5. HTP-3 intensity was adjusted differently for the two genotypes due to fainter staining in *duo-1* spreads (see Fig S6). Scale bar = 10µm. *D)* Immunofluoresence images of HTP-3 and crossover-associated factor COSA-1, showing that COSA-1 foci are present in the *duo-1* mutant and colocalize with SC aggregates. Scale bar = 10µm. *E)* Quantification showing reduced and varied numbers of COSA-1 foci in *duo-1* mutants.

Removal of RAD-51 foci during wild-type meiosis reflects progression of DSB repair to later stages. In principle, RAD-51 hyperaccumulation and persistent high numbers of foci in *duo-1* germ lines could reflect failure of early DSB repair intermediates to progress to later stages, continued induction of DSBs throughout pachytene, and/or continued loading of RAD-51 on existing intermediates when meiotic structures become dismantled. We addressed the issue of repair progression by co-staining for RAD-51 and meiotic recombination factor MSH-5, which marks post-strand-exchange repair intermediates (26, 30, 47) (Fig 3C). Despite persistence of RAD-51 foci in *duo-1* meiotic prophase nuclei, MSH-5 foci were also detected, suggesting that a subset of DSBs do indeed progress to form later intermediates. However, RAD-51 foci vastly outnumbered MSH-5 foci in the *duo-1* mutant, in striking contrast to the situation in wild-type meiosis.

Finally, we assessed localization of COSA-1, a cyclin-like protein that accumulates during the late pachytene stage in wild-type meiosis at the subset of DSB repair sites selected to become crossovers (COs) (30) (Fig 3D-E, S6). Bright COSA-1 foci were indeed detected in late-pachytene region nuclei in the *duo-1* mutant, albeit numbers of foci were variable and reduced relative to the 6 foci (1 per chromosome pair) reliably observed in wild-type meiocytes. Notably, COSA-1 foci in the *duo-1* mutant correlated in numbers with and were consistently colocalized with SC protein polycomplexes. As polycomplexes are still observed in the *duo-1; spo-11* double mutant, the formation of these polycomplexes is not dependent upon the presence of recombination intermediates; however, SC components may be preferentially retained where intermediates form. Colocalization of MSH-5 and COSA-1 in late prophase foci in *duo-1* mutants (Fig S6) is consistent with the possibility that such foci may represent sites of intermediates that had progressed to late CO-fated stages of repair. However, given the impairment of homolog pairing in *duo-1* mutants, we infer that if these MSH-5 and COSA-1 foci do represent repair intermediates, such intermediates may be occurring between sister chromatids.

### *duo-1* mutants temporarily engage, then deactivate, the crossover assurance checkpoint

The protein kinase CHK-2 is a key regulator of meiotic progression in the *C. elegans* germ line. CHK-2 becomes activated at the onset of meiotic prophase and is an essential driver of early prophase nuclear reorganization, homolog pairing, proper SC assembly, and induction of programmed DSBs (48, 49). Further, exit from the early pachytene stage is coupled to cessation of CHK-2 activity, reflecting satisfaction of a “CO assurance checkpoint” that senses whether all chromosomes have successfully formed potential CO intermediates (17, 18, 50–52). Failure to satisfy this checkpoint prolongs CHK-2 activity, which can be visualized in mutants with impaired synapsis and/or recombination as an extended zone of nuclei exhibiting markers of CHK-2 activity, such as phosphorylation of nuclear envelope protein SUN-1 at Serine 8 (SUN-1 pS8) (53).

Immunostaining for SUN-1 pS8 shows that *duo-1* mutants exhibit a significant extension of the CHK-2 active zone, indicating that nuclei are at least temporarily able to detect deficits in homolog engagement (Fig S7). However, the degree of the SUN-1 pS8 zone extension is less than is observed in mutants lacking SYP proteins or in other mutants exhibiting similar degrees of RAD-51 hyperaccumulation (18, 50). As meiotic axis proteins HTP-3 and HIM-3 are required for operation of the CO assurance checkpoint (18, 51), we infer that progressive loss of axis integrity during meiotic progression in *duo-*1 mutant germ lines results in eventual inability to sustain checkpoint activation.

### Temporally-controlled DUO-1 depletion reveals separable roles for DUO-1 in SC assembly, in SC maintenance, and in maintenance of chromosome compaction during late prophase

In principle, the progressive deterioration of meiotic chromosome structure and the severe late prophase decompaction phenotype observed in *duo-1* null mutants might represent either: a) downstream consequences of loss of a key DUO-1 function during (or prior to) early prophase, or b) a continued requirement for DUO-1 function during prophase progression. To distinguish between these possibilities, we used the auxin-inducible degron (AID) system (54, 55) to enable temporally-controlled degradation of AID-tagged DUO-1 protein (Fig 4). By assessing the outcomes of acute and longer-term DUO-1 depletion, we uncovered multiple distinct roles for DUO-1 during meiotic prophase progression.

**Figure 4.**
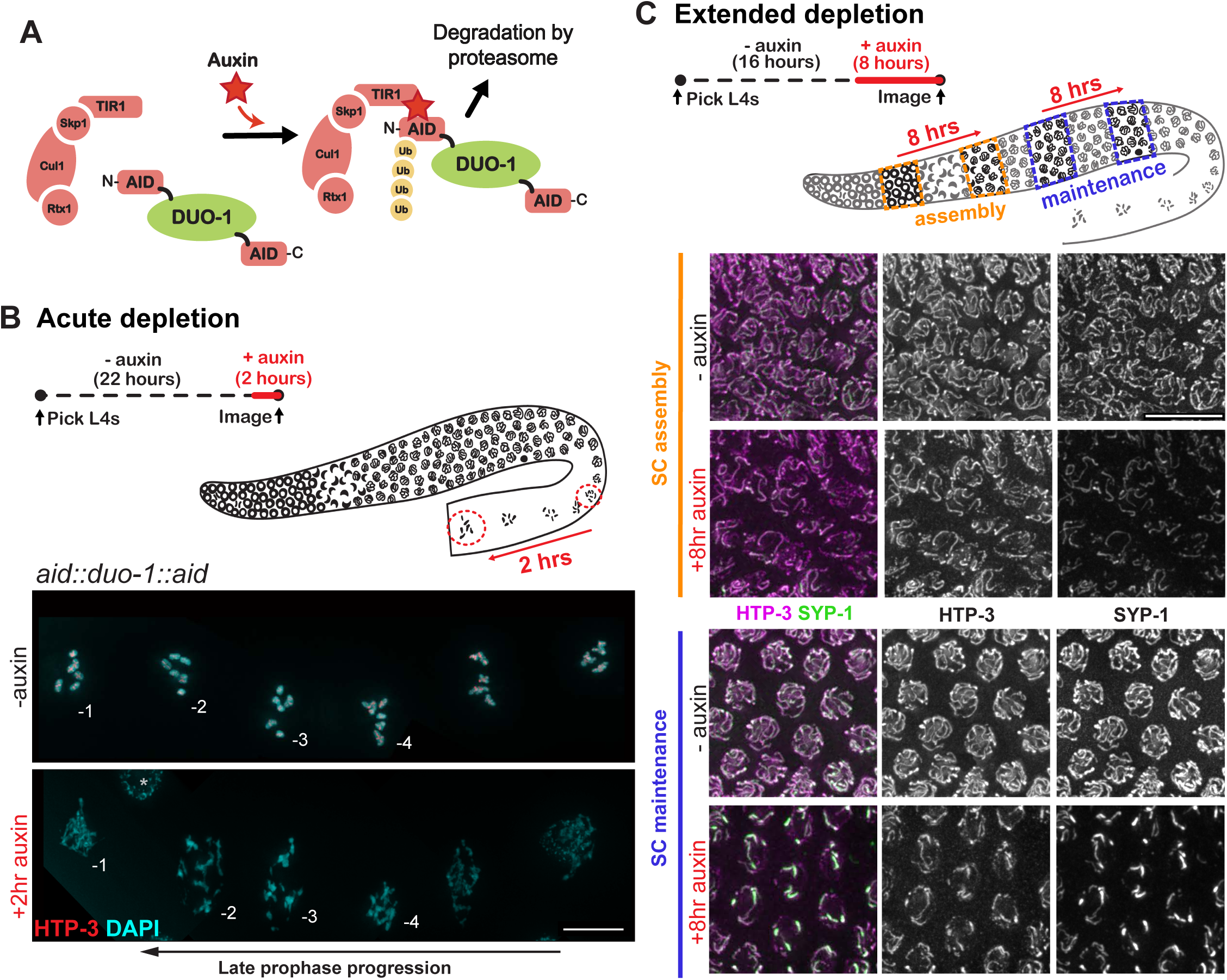
Auxin-induced DUO-1 depletion reveals separable roles for DUO-1 in SC assembly, in SC maintenance, and in maintenance of chromosome compaction during late prophase. *A)* Schematic of the auxin inducible degron (AID) system employed to temporally deplete DUO-1. *B)* Acute depletion of DUO-1 recapitulates late prophase chromatin compaction phenotypes seen in *duo-1* mutants. Top: schematics depicting acute auxin depletion treatment and progression of nuclei of interest in the germ line during the two-hour treatment window. Bottom: images of proximal diakinesis oocytes from *aid::duo-1::aid* worms with and without addition of auxin for 2 hr, showing decondensed chromatin following acute DUO-1 depletion (asterisk indicates a somatic sheath-cell nucleus). Scale bar = 10µm. *C)* 8-hr auxin treatment reveals separate roles for DUO-1 in assembly and in maintenance of SCs. Top: schematics depicting 8-hr auxin treatment and progression of nuclei of interest to assess effects on SC assembly in nuclei entering meiotic prophase during the treatment (orange) and on SC maintenance in nuclei where the SC had fully assembled prior to DUO-1 depletion (blue). Bottom: images of HTP-3 and SYP-1 in nuclei from the corresponding end-point regions indicated in the schematic. Scale bar = 10µm.

Remarkably, we found that an acute 2 hour auxin exposure was sufficient to recapitulate the severe chromosome decompaction phenotype observed in diakinesis-stage oocyte nuclei in *duo-1* null mutants (Fig 4B). Following a 2-hour auxin treatment, DAPI-stained chromatin in diakinesis-stage oocytes appeared decondensed, stringy and/or diffuse, and no evidence of HTP-3-marked axial structures was detected. Importantly, the chromosomes in late diakinesis oocytes at the end of this acute auxin treatment (*i.e.* in the −1 and −2 positions relative to the spermatheca) would already have achieved the tightly-compacted diakinesis bivalent structure prior to the time when DUO-1 was depleted. Thus, this rapid loss of chromosome organization upon DUO-1 depletion indicates that DUO-1 plays an important role during the diakinesis stage to maintain the compact condensed structure of late prophase meiotic bivalents.

To evaluate the effects of DUO-1 depletion during earlier stages of meiotic prophase, we conducted an 8-hour auxin treatment followed by fixation and immunostaining for HTP-3 and SYP-1 (Fig 4C). Since nuclei travel forward in the germ line by about 8 rows during the 8 hour time frame of this treatment (42, 43), this approach enabled us to evaluate the status of SCs in both: a) cohorts of early prophase nuclei that would not yet have assembled their SCs prior to DUO-1 depletion, and b) cohorts of nuclei in the late pachytene region that would have already completed SC assembly prior to DUO-1 depletion. HTP-3 and SYP-1 immunostaining showed that SC assembly was incomplete in early prophase nuclei following 8-hour auxin treatment, confirming a role for DUO-1 in SC assembly inferred from analysis of the null mutant. Moreover, we observed reduced and aberrant HTP-3 and SYP-1 signals in nuclei in the late pachytene region, and by entry to diplotene, we observed a near-complete breakdown of the SC into large polycomplexes (Fig S8), demonstrating a role for DUO-1 in maintenance of axis and SC organization and integrity even after these structures have been fully assembled.

Together, these data demonstrate an ongoing requirement for DUO-1 throughout meiotic prophase progression, reflecting multiple distinct roles for DUO-1: a) in promoting proper axis/SC assembly in early prophase, b) in maintaining axis/SC structure during the late pachytene stage, and c) in promoting and maintaining chromosome compaction at the end of meiotic prophase. More broadly, this work reveals a previously unappreciated requirement for active maintenance of meiosis-specific chromosome structures and meiotic chromosome architecture throughout meiotic prophase.

### Tagging DUO-1 for localization and proximity labeling in germ cell nuclei

To visualize localization of DUO-1 in the germ line, we used CRISPR-Cas9 genome editing to create strains expressing either 3xFLAG::DUO-1 or DUO-1::GFP from the endogenous *duo-1* locus. Consistent with the meiotic roles of DUO-1 described above, immunofluorescence and/or live imaging indicate that 3xFLAG::DUO-1 and DUO-1::GFP are enriched in nuclei throughout the germ line and increase in abundance during late prophase progression (Fig 5A-B). Further, DUO-1 is detected predominantly in the nucleoplasm (Fig 5C). However, a nuclear spreading protocol that releases nucleoplasmic protein pools reveals that a subset of DUO-1 colocalizes with the chromosome axes (Fig 5D); this axis-associated DUO-1 signal is most readily detected during the late pachytene/ early diplotene stage.

**Figure 5.**
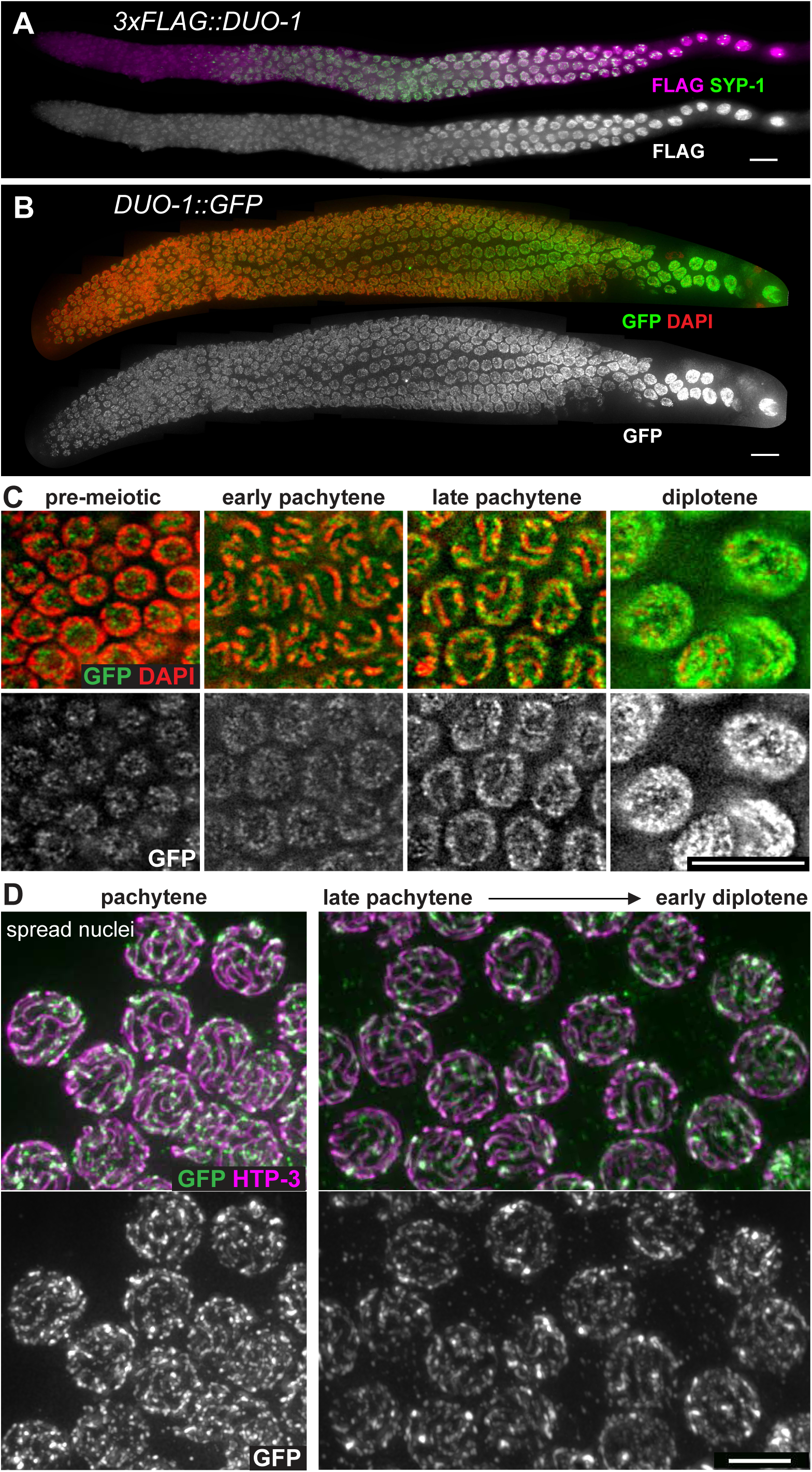
DUO-1 localizes to germline nuclei throughout meiotic prophase. Immunolocalization of *A)* 3xFLAG::DUO-1 and *B)* DUO-1::GFP in whole mount gonads, showing enrichment in germline nuclei that becomes more pronounced in late prophase. Scale bar = 10µm. *C)* Single z-slice images showing that DUO-1::GFP is localized primarily in the nucleoplasm. Scale bar = 10µm. D) Late pachytene and early diplotene nuclei from nuclear spread preparations that release soluble protein pools, showing DUO-1::GFP localizing to meiotic chromosome axes. Scale bar = 5µm.

To identify proteins that associate with DUO-1 in germ cell nuclei, we employed a modular proximity-labelling approach involving tissue-specific expression of the TurboID biotin ligase fused to a GFP nanobody (56) to couple the biotin ligase activity to DUO-1::GFP. To improve specificity and ensure relevance of our controls, we introduced a nuclear localization signal (NLS) into the nanobody-TurboID coding sequence of a strain expressing the nanobody-TurboID fusion protein specifically in the germ line. (Fig S9; Methods).

Our analysis of proteins preferentially biotinylated in worms co-expressing nanobody-NLS-TurboID and DUO-1::GFP in their germ lines is presented in Fig 6A and Fig S9. DUO-1 itself was strongly enriched in experimental vs. control samples, providing internal validation. Further, known meiotic axes components (cohesion subunits HIM-1 and SMC-3 and HORMAD protein HIM-3) were significantly overrepresented among the 19 additional proteins that exhibited both >4-fold increased abundance in experimental samples and p-values < 10^-2^. This overrepresentation of meiotic axis proteins is consistent with the demonstrated role for DUO-1 in maintaining axis integrity and sister chromatid cohesion. While the numbers of peptides detected for these proteins individually were not high enough for them to be flagged as high-confidence interactors, this may reflect the fact that the bulk of DUO-1 is in the nucleoplasm, where it likely interacts with other partners.

**Figure 6.**
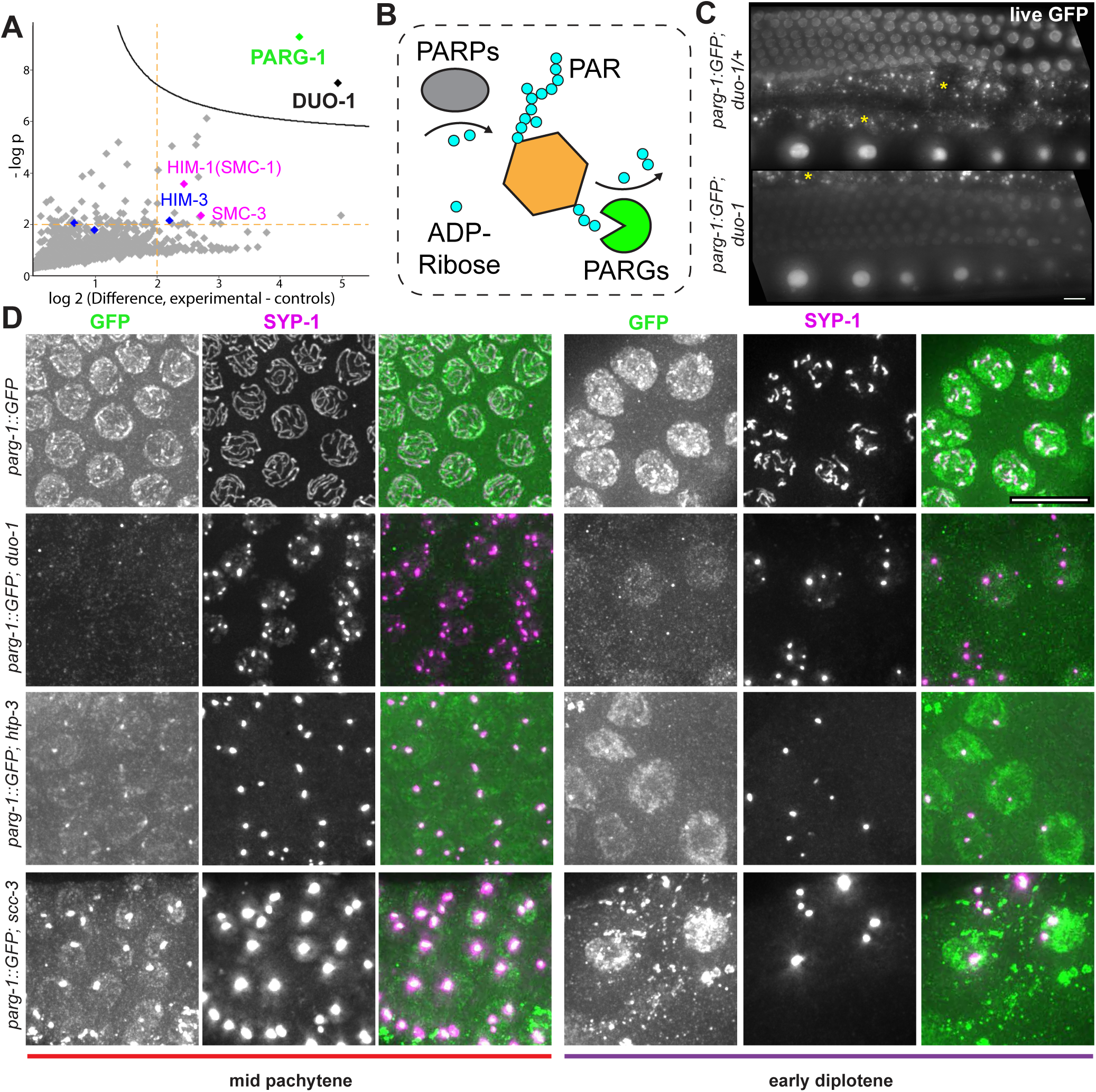
TurboID and protein visualization reveal a connection between DUO-1 and PARG-1. *A)* One-sided volcano plot of data from DUO-1:GFP TurboID analysis, where each point represents a protein identified by LC-MS from biotinylated proteins captured by streptavidin pull down and purification. x-axis shows log_2_ fold difference in abundance between experimental (*GFPnanobody ::NLS::TurboID* + *duo-1::GFP*) and two sets of controls (*duo-1::GFP* only and *GFPnanobody::NLS::TurboID only*), and y-axis shows −log 10 of the p value. Stringent significance curve in the top right of plot was generated in Perseus (FDR = 0.05, s0 = 0.1). Blue points correspond to meiotic HORMAD proteins and pink points correspond to cohesin components. See Methods and Supplement for full details and strain genotypes. *B)* Schematic of Poly-ADP-Ribosylation (PARylation) cycle, showing the interplay of PARPs, which add ADP-ribose to protein targets, and PARGs, which dismantle PAR and remove ADP-ribose from protein targets. *C)* Max projection images of *PARG-1::GFP* fluorescence in live worms, showing reduced *PARG-1:GFP* fluorescence in germline nuclei in a *duo-1* mutant background (yellow asterisk: background autofluorescence of gut). Scale bar = 10µm. *D)* Immunofluorescence images of *PARG-1::GFP* in whole mount germ lines from worms of indicated genotypes, showing reduced nuclear and chromosomal localization and reduced enrichment in polycomplexes in the *duo-1* mutant. All images are from worms dissected and stained in tandem with identical imaging parameters and post-processing adjustments (resulting in post-processing pixel oversaturation in some images). Scale bar = 10µm.

Indeed, our nanobody-TurboID analysis identified a nucleoplasmic protein, Poly(ADP-ribose) Glycohydrolase 1 (PARG-1), as a high-confidence DUO-1 interacting partner (Figure 6A and Fig S9). PARG-1 is the primary PAR glycohydrolase in the *C. elegans* germ line (57), and like DUO-1, it exhibits nucleoplasmic localization throughout the meiotic germ line as well as colocalization with axis/SC components that becomes most prominent in late pachytene and early diplotene nuclei (57) (Fig 6C-D). While identification of PARG-1 as a major DUO-1 interacting partner might not have been anticipated based the relatively mild impairment of meiosis observed in *parg-1* single mutants (57, 58), DUO-1 was independently identified as a candidate PARG-1 interacting protein in an IP/MS experiment targeting PARG-1::GFP (Fig S9).

### PARG-1::GFP localization is impaired in duo-1 mutants

We used live imaging and immunofluorescence microscopy to visualize PARG-1::GFP in the *duo-1* mutant (Fig. 6C-D; Fig S9). Nuclear-enriched PARG-1::GFP fluorescence was readily visible in germline nuclei of live *WT* and *duo-1/+* control worms, and immunofluorescence images showed colocalization of a subset of PARG-1::GFP with SCs and with SYP-1-enriched domains in late pachytene and diplotene nuclei. In contrast, both live PARG-1::GFP fluorescence and immunofluorescence detection of PARG-1::GFP colocalization with SYP-1 (in polycomplexes) were diminished in *duo-1* mutant germ lines. Reduced detection of PARG-1::GFP associated with SYP-1 in the *duo-1* mutants is not simply a consequence of SC proteins being concentrated in polycomplexes (rather than assembled SCs), as we readily detected co-enrichment of PARG-1::GFP with SYP-1 in the polycomplexes that form in mutants lacking axis components HTP-3 or SCC-3 (Fig 6D).

## Discussion

### DUO-1 promotes active maintenance of meiotic chromosome architecture

After more than 30 years of generating and analyzing *C. elegans* meiotic mutants, it was refreshing to see a new meiotic phenotype emerge from our recent screening efforts. The *duo-1* mutants stood out immediately based on the highly decompacted state of chromosomes in *duo-1* diakinesis oocytes. This contrasts both with the compact univalents observed in the vast majority of meiotic mutants – those impaired for homolog pairing, axis/SC assembly, formation of DSBs and/or repair of DSBs as crossovers (5, 22, 29, 32, 39, 48) – and with the fragmented/aggregated chromosomes observed in some mutants defective for DNA repair (46, 59). Indeed, a similar diakinesis phenotype has been reported previously only for conditional depletion of condensin II (60; see below). The unique phenotype of *duo-1* mutants, coupled with the rapid deterioration of chromosome structure upon DUO-1 depletion, have provided new insights into meiotic prophase chromosome dynamics.

Specifically, our data indicate that meiotic chromosome structures are less stable than we had previously appreciated but instead require active maintenance throughout meiotic prophase. Chromosomes are apparently vulnerable to being dismantled on a moment’s notice and consequently require continuous protection to maintain structural stability. DUO-1 has an essential role in conferring this protection.

Our examination of *duo-1* mutant phenotypes has revealed several vulnerable features that require DUO-1 for protection. First, we have demonstrated that the integrity of meiotic chromosome axes requires the protection of DUO-1, and several lines of evidence suggest that meiosis-specific cohesin complexes are the vulnerable targets. REC-8 cohesin is present in the SCs that initially form in *duo-1* mutants but becomes depleted as chromosome axes and SCs disassemble and SYP-1 and HORMAD proteins become concentrated into polycomplexes, concomitant with loss of sister chromatid cohesion. Cohesin complex impairment can likewise explain the hyper-accumulation of RAD-51-marked early DSB repair intermediates, as similar accumulation of RAD-51 foci occurs in *rec-8* mutants (35, 46). Moreover, as REC-8 is not required to establish or maintain meiotic chromosome axes (6, 10, 44, 61), we can also infer that not only REC-8 cohesin, but rather both types of meiosis-specific cohesin complexes require DUO-1-mediated protection. Further, we detected a >6-fold enrichment of the SMC-1 component shared by both meiotic cohesin complexes (44) in our DUO-1 TurboID proximity labeling experiments, consistent with cohesin complexes being DUO-1 clients.

It is well established that cohesin complexes are targets of cell cycle-regulated degradation to enable chromosome segregation during both mitotic and meiotic cell divisions (62, 63). Moreover, it is well recognized that there is a need for temporary localized protection of a subset of cohesin complexes during meiosis in order to enable the two-step release of sister chromatid cohesion during the two meiotic divisions (64, 65). Our work shows that the need for active protection of cohesin complexes starts much earlier and is important for maintaining chromosome axis structure in the context of the SC during the pachytene stage. We speculate that this proposed cohesin-protection function conferred by DUO-1 acts in parallel with the ongoing SCC-2/ Nipped B-dependent dynamic turnover of cohesin complexes that has been observed during the pachytene stage in both *C. elegans* and *Drosophila* (6, 66, 67). Cohesin protection can’t be the whole story, however. In contrast to the dramatic chromosome decompaction phenotype observed in *duo-1* mutants at the end of meiotic prophase, absence of cohesin complexes in *C. elegans* meiotic germ cells results in compact individual chromatids and chromosome fragments in diakinesis oocytes (44). The *duo-1* diakinesis decompaction phenotype is more reminiscent of effects observed upon depletion of condensin II during meiosis (68), raising the possibility that the condensin II complex might be a late prophase beneficiary of DUO-1-mediated protection. Interestingly, condensin II loads onto meiotic chromosomes after SC disassembly to mediate chromosome compaction during the diplotene/diakinesis stages. This timing corresponds to the window during meiotic progression when chromosomes are acutely vulnerable to rapid loss of structure upon DUO-1 depletion.

### Deubiquitinating enzymes and meiotic chromosome structure

Based on the identity of DUO-1 as an ortholog of conserved deubiquitinating enzymes, it is natural to envision that DUO-1 may confer protection of chromosome architecture by antagonizing ubiquitin-mediated protein degradation. There is a growing body of literature implicating protein ubiquitination and ubiquitin-mediated proteolysis as important regulators of meiotic functions. The proteasome itself has been reported to localize to nuclei and/or on chromosome axes during meiotic prophase in multiple organisms, including mice, yeast and *C. elegans* (*e.g.* refs. 69–72) and inhibition or loss of proteasome function in these contexts impairs morphogenesis of chromosome axes and/or leads to persistence of non-productive early assemblages of SC components (69, 72). There have also been numerous reports of predicted RING-finger E3 ligases or predicted components or regulators of SCF-like E3 ubiquitin ligases affecting various aspects of the meiotic prophase program. For example, the *C.elegans* SCF^PROM-1^ complex plays key roles in promoting meiotic prophase entry and homolog pairing (73–75). Further, the *C. elegans* CSN/COP9 signalosome has been implicated in promoting SC assembly by regulating Cullin activity to antagonize aggregation of SC subunits (76), However, there are only a few reports of meiotic prophase roles for deubiquitinating enzymes that counteract the activities of ubiquitin ligases by removing ubiquitin from proteins.

Paralleling our finding that *C. elegans* DUO-1 is required to maintain assembled meiotic axes and SCs, a recent report by Lake *et al.* (2024) demonstrated a role for a different class of deubiqutinating enzyme, ubiquitin-specific protease Usp7, in maintaining SCs during *Drosophila* oocyte meiosis (77). However, in the *Drosophila* study, it was not possible to discern whether SC instability reflected an inherent vulnerability in the assembled SC structures or an ongoing requirement for active maintenance. Our temporal depletion of DUO-1 by AID allowed us to infer a continued need for active maintenance, by showing that even in nuclei with fully functional assembled SCs, the structure cannot be maintained when DUO-1 is depleted. There may also be distinctions between the underlying points of structural vulnerability of the SCs in these two systems, as our data point to meiotic cohesin complexes as an entity in need of protection in *C. elegans*, whereas no loss of cohesion was detected following Usp7 knockdown in *Drosophila*, suggesting that some other protein(s) may be the weak link in *Drosophila* oocytes. (However, the authors noted that cohesion was assessed only at centromeres, so potential loss of cohesion outside of centromeric regions could have been missed.) Together, these complementary studies suggest that promoting maintenance of SC structure by antagonizing protein ubiquitination may be a conserved feature of the meiotic program, albeit the specific proteins or structures in need of protection may vary.

### The role of DUO-1/PARG-1 Interaction

During the course of this work, we uncovered the Poly(ADP-ribose) glycohydrolase PARG-1 as a high-confidence DUO-1 interacting partner and demonstrated that DUO-1 is required for normal abundance and localization of PARG-1 in germ cell nuclei. However, we know that PARG-1 cannot be an essential co-factor for DUO, as *parg-1* single mutants exhibit only a very modest impairment of meiosis and produce mostly viable embryos (57), indicating the presence of functional DUO-1. What, then, is the functional significance of this DUO-1/PARG-1 partnership?

Our consideration of potential functional connections is informed by prior research that had: 1) demonstrated interactions between PARG-1 and several axis/SC components, 2) identified (through double and triple mutant analyses) roles for PARG-1 in augmenting and coordinating processes of meiotic DSB formation and repair, and 3) demonstrated that catalytically-inactive PARG-1 protein is at least partially functional in promoting meiotic recombination (57, 58). We speculate that DUO-1 might serve to facilitate interaction of PARG-1 with axis/SC components and/or support PARG-1 in its recombination-promoting roles. However, such potential contributions of DUO-1 to PARG-1 function would be difficult to discern given the reduced detection of PARG-1 and the profound loss of chromosome architecture in *duo-1* mutants. Alternatively, or in addition, PARG-1 could have a (non-essential) role in facilitating association of DUO-1 with chromosome axis.

### Concluding remarks

Together, our data reveal that meiosis-specific chromosome structures and meiotic chromosome architecture require active maintenance throughout meiotic prophase. *C. elegans* DUO-1 plays an essential role in this maintenance, conferring protection at multiple junctures during meiotic progression to prevent dismantling of chromosome organizational features necessary for transmission of intact, functional genomes to the next generation.

## Materials & Methods

### C. elegans genetics and genome engineering

Worms were cultured using standard methods (78) at 20°C. A list of strains used is provided in the SI Appendix. Established approaches were used to re-create mutations identified by genetic screening, to create a STOP-IN null allele of *duo-1*, and to introduce endogenous tags at the *duo-1* locus (28, 79, 80) (see Supplemental Methods for sequence and design information).

*duo-1* mutations identified by genetic screening:

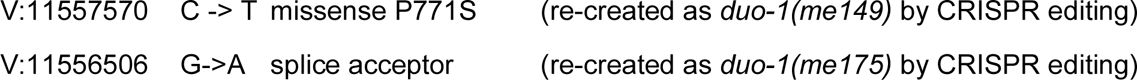

### Immunofluorescence

Immunofluorescence experiments involving whole mount gonads or nuclear spread preparations were conducted as in (9, 15, 81), with modifications, using adult hermaphrodites 24 - 28 hours post-L4 stage. Antibodies used, details of procedures, and image acquisition and processing are provided in SI.

### FISH

FISH experiments were conducted as in (82) using adult hermaphrodites 24 - 28 hours post-L4 stage.

### Auxin-inducible degradation (AID) Experiments

For AID experiments, worms were cultured using standard methods and picked to new plates as late L4s. Animals were then transferred to plates containing 2mM indole-3 acetic acid (IAA) as in (55), after 16 hours (for 8 hour auxin treatments) or after 22 hours (for acute 2 hour auxin treatments). All worms were dissected at 24 hours post-L4 and fixed for immunofluorescence. Controls were cultured on standard NGM plates.

### Imaging

Images of whole-mount gonads or nuclear spread preparations were acquired on a DeltaVision OMX Blaze microscope with a 100 x 1.4 NA widefield objective and z-stacks spaced at 200nm, deconvolved and registration corrected with SoftWoRx software and, where applicable, stitched together using the ‘Grid/Collection stitching’ FIJI plugin (83).

### Image quantification

For FISH experiments, foci were quantified and assigned to individual nuclei in whole-mount gonads using the 3D Maxima Finder plugin in FIJI (84). Details on 3D maxima-finder quantification, assignment of foci to nuclei, and zone segmentation can be found in the SI Appendix.

For quantification of RAD-51 foci, max projected images of Z-stacks of stitched whole-mount germ lines were straightened using the ‘straighten’ function in FIJI. The length of each germ line was measured from distal tip to the region of the proximal germ line where number of nuclei per row decreased to 1-2. Total RAD-51 foci were then quantified using 3D maxima finder, which outputs xy locations of foci as well as peak intensities. Positions of foci along the distal-proximal axis of the germ line were normalized to germline length and plotted in GraphPad Prism.

### LC/MS Sample preparation and data analysis for TurboID

Nine biotinylated protein samples (three replicates per condition) were digested on streptavidin-coated beads and prepared for LC-MS/MS analysis as described in SI Appendix before ionization and analysis using an Orbitrap Exploris 480 mass spectrometer at the Stanford University Mass Spectromery (SUMS) core. The .RAW data were analyzed using Byonic v5.2.5 (Protein Metrics, Cupertino, CA) to identify peptides and infer proteins. See SI Appendix for full details.

Data were further processed and visualized using the Perseus software package (85). Specifically, data were filtered to remove contaminants, log2 transformed, and trimmed to remove rows with insufficient data (>3 NaN values). To determine Fold Change, imputation was used to add a small constant (0.01) to all values in the dataset. Data were plotted as a one-sided volcano plot using this software for Figure 6A.

### Statistical analyses

Statistical analyses were performed using GraphPad Prism. Specific tests are described in the figure legend of the figures that they appear in.

## Acknowledgments

We thank C. Akerib and R. Yokoo and members of the Villeneuve lab for genetic screening efforts and for discussions and comments on the manuscript. We thank C. Uebel for germ line schematics in figure 4. We thank M. Zetka, A. Dernburg, D. Libuda, S. Wignall and V. Jantsch for antibodies and the Caenorhabditis Genetics Center (funded by NIH Office of Research Infrastructure Programs P40 OD010440) for strains. TurboID proteomics work was supported by the Vincent Coates Foundation Mass Spectrometry Laboratory, Stanford University Mass Spectrometry (RRID:SCR_017801) utilizing the Thermo Exploris 480 nanoLC/MS system (RRID:SCR_022215) and supported in part by NIH P30 CA124435 utilizing the Stanford Cancer Institute Proteomics/Mass Spectrometry Shared Resource. Proteomics analyses in Fig S9D were performed by the Mass Spectrometry Facility at Max Perutz Labs using the VBCF instrument pool. This work was supported by an American Cancer Society Research Professor Award (RP-15-209-01-DDC) and NIH grant R35GM126964 to AMV, by Czech Science Foundation award GA23-04918S to NS, and by NIH grant 1S10OD01227601 from the NCRR to the Stanford Cell Sciences Imaging Facility (RRID:SCR_017787). LGS was supported by NIH Training Grant T32GM007790 and CPC by a Stanford Medicine Dean’s Postdoctoral Fellowship.

## Notes

### Competing Interest Statement

The authors have declared no competing interest.

