## Supplementary Information for "Active maintenance of meiosis-specific chromosome structures in *C. elegans* by the deubiquitinase DUO-1"

Includes:

**Supplemental Methods**

**Supplemental References**

**Supplemental Figure Legends**

**Supplemental Figures S1- S9**

#### Supplemental Materials and Methods

##### *C. elegans* strains

The following *C. elegans* strains were used in this paper and are available from AMV upon request:

**AV1206** *duo-1(me149[P771S])/ tmC12 [egl-9(tmIs1194)] V*

**AV1281** *duo-1(me175[splice acceptor])/ tmC12 [egl-9(tmIs1194)] V*

**AV1306** *duo-1(me186[G4 STOP-IN]) / tmC12 [egl-9(tmIs1194)] V*

**DAM1284** *vieSi142[pAD861; Ppie-1::vhhGFP4::HA::TurboID::operon-linker::mCherry::his-11::tbb-2 3'UTR; cb unc-119(+)] II; unc-119(ed3) III*

**AV1308** *vieSi142[pAD861; Ppie-1::vhhGFP4::HA::TurboID::operon-linker::mCherry::his-11::tbb-2 3'UTR; cb unc-119(+)] II; unc-119(ed3) III; duo-1(me187[duo-1::flexlinker::eGFP])* V

**AV1379** *vieSi142[pAD861; Ppie-1::vhhGFP4::NLS::HA::TurboID::operonlinker::mCherry:: his-11::tbb-2 3'UTR; cb unc-119(+)] II; unc-119(ed3) III*

**AV1380** *vieSi142[pAD861; Ppie-1::vhhGFP4::NLS::HA::TurboID::operonlinker::mCherry:: his-11::tbb-2 3'UTR; cb unc-119(+)] II; unc-119(ed3) III; duo-(me187[duo-1::flexlinker::eGFP])V;*

**AV1227** *duo-1(me154[3xFLAG::duo-1])*

**AV1289** *gld-1::tir1::mRuby II; duo-1(me179[aid::duo-1::aid]) V*

**AV1422** *gld1:tir1:mRuby II; him-6(jf93[him6:3xHA]) msh-5::V5 IV; duo-1(me179[aid::duo-1::aid]) V*

**AV1399** *spo-11(me44)/ tmC5 [F36H1.3(tmIs1220)] IV; duo-1(me186)/ tmC12 [egl-9(tmIs1194)] V*

**NSV24** *parg-1[ddr2(parg-1::GFP)] IV*

**AV1398** *cosa-1(ddr12[ollas::cosa-1]) III; parg-1[ddr2(parg-1::GFP)] IV; duo-1(me186) / tmC12 [egl-9(tmIs1194)] V*

**AV1417** *parg-1[ddr2(parg-1::GFP)] IV; scc-3(ku263) / tmC12 [egl-9(tmIs1194)] V*

**AV1423** *htp-3(y428) ccIs4251 I/hT2[bli-4(e937)let-?(q782) qIs48] (I,III); parg-1[ddr2(parg-1::GFP)] IV*

##### CRISPR-Cas9 Gene Editing

*duo-1(me149)* was generated using CRISPR-Cas9 gene editing as described in (1), using Cas9 protein (0.25µg/µL), tracrRNA (0.02µg/µL, and crRNA (0.02µg/µL) (5'- TTCTTTCTCAATGAAGT TGT[AGG] - 3'), dpy-10(cn64) (40ng/µL) (2) . A ssODN (0.11µg/µL) was used as the repair template (5' – CACCAAGAAAAGGAGCTATATATTTGAAGAATACAAATATGCATTATGAGTCT ACAACTTCATTGAGAAAGAATTCACCAGAAAGGTCGCCAATGAAAAAAGA - 3'). F1 rollers were screened by PCR using the following primers (5'- AATTTACATTAAGCACCTTCAGTCGT AC - 3' and 5'- CAAGAAGAACGAATTAATGTTC AAAAAGTACTG - 3') followed by a Acc1 restriction enzyme digest. The edit was confirmed by Sanger sequencing.

*duo-1(me175)* was generated using CRISPR-Cas9 gene editing as previously described, using Cas9 protein (0.25µg/µL), tracrRNA (0.02µg/µL, and crRNA (0.02µg/µL) (5'-GATTCACAGGGATGGATGTT[CGG]-3'), dpy-10(cn64) (40ng/µL). A ssODN (0.11µg/µL) was used as the repair template (5' – TAGACGACACATTTATGATTCACAAGGATGGATGTTCAG TGCAAAAACTGGTTTCGTCGTTTGTTAAATGGTC - 3'). F1 rollers were screened by PCR using the following primers (5'- ATCCAGCAAG GGCTTTCAAG - 3' and 5'- GAAAAAGGAGGT CCTTGTCC - 3') followed by a BtsIMutI restriction enzyme digest. The edit was confirmed by Sanger sequencing.

*duo-1(me186)* was generated using CRISPR-Cas9 gene editing as previously described, using Cas9 protein (0.25µg/µL), tracrRNA (0.02µg/µL, and crRNA (0.02µg/µL) (5'- TTAGATGACCGA AGGGGAAGTTTGTCCAGAGCAGAGGTGACTAAGTGATAAGCTAGCAATTTA [TGG]-3'), dpy-10(cn64) (40ng/µL). A ssODN (0.11µg/µL) was used as the repair template with 35 bp homology arms flanking a 43bp universal STOP-in cassette (8) (5' – GCATATAATTCAAAAGGA TTTTTAGATGACCGAAGGGGAAGTTTGTCCAGAGCAGAGGTGACTAAGTGATAAGCTAGCA ATTTATGGTTGATGTCTCAATTGTCGGTTCGGAT - 3'). F1 rollers were screened by PCR using the following primers (5'- TCGGTGTGATGCATCATTATTGC -3' and 5'- ATGAAGTGAAG ATCCATCGAATTTTAGTC - 3'). Successful insertion resulted in a larger fragment and was verified by sequencing.

*duo-1(me154[3xFLAG::DUO-1])* was generated using CRISPR-Cas9 gene editing as previously described, using Cas9 protein (0.25µg/µL), tracrRNA (0.02µg/µL, and crRNA (0.02µg/µL) (5'-AAGAATTTATGGTTGATGTC [CGG]-3'), dpy-10(cn64) (40ng/µL). A ssODN (0.11µg/µL) was used as the repair template (5' – TATAAATTAAGCATATAATTCAAAAGGATTTTTAGATGGG ATCGGACTATAAAGATCACGACGGAGATTACAAGGACCATGATATCGACTACAAGGACGAC GACGACAAGGGAGGAGGCTCAGGAACCGAAGAATTTATGGTTGATGTCTCAATTGTCGG-3'). F1 rollers were screened by PCR using the following primers (5'- TCGGTGTGATGCATCA TTATTGC -3' and 5'- ATGAAGTGAAGATCCATCGAATTTTAGTC - 3'). Successful insertion resulted in a larger fragment and was verified by sequencing.

*duo-1(me179[AID::DUO-1::AID])* was generated using a two-step CRISPR-Cas9 gene editing as previously described, to add a degron to first the N terminus, then the C terminus. For the N-terminus, we used Cas9 protein (0.25µg/µL), tracrRNA (0.02µg/µL, and crRNA (0.02µg/µL) (5'-AAGAATTTATGGTTGATGTC [CGG]-3'), dpy-10(cn64) (40ng/µL). A ssODN (0.11µg/µL) was used as the repair template (5' – GCATATAATTCAAAAGGATTTTTAGATGCCTAAAGATCCA GCCAAACCTCCGGCCAAGGCACAAGTTGTGGGATGGCCACCGGTGAGATCATACCGGAA GAACGTGATGGTTTCCTGCCAAAAATCAAGCGGTGGCCCGGAGGCGGCGGCGTTCGTGA AGGGAGGCTCAGGAACCGAAGAATTTATGGTTGATGTCTCAA- 3'). F1 rollers were screened by PCR using the following primers (5' – TCGGTGTGATGCATCATTATTGC -3' and 5'-ATGAAGTGAAGATCCATCGAATTTTAGTC - 3'). Successful insertion resulted in a larger fragment and was verified by sequencing. For C terminus degron insertion, the same schema was used with crRNA (0.02µg/µL) (5'- CATGGAAAAATAATTTCTAC [TGG]-3') and ssODN (0.11µg/µL) was used as the repair template (5' –GTCCGGATATCTTATGTTTTATGAACTCCA

GGGAGGCCCTAAAGATCCAGCCAAACCTCCGGCCAAGGCACAAGTTGTGGATGGCCAC CGGTGAGATCATACCGGAAGAACGTGATGGTTTCCTGCCAAAAATCAAGCGGTGGCCCGG AGGCGGCGGCGTTCGTGAAGTAGAAATTATTTTTCCATGTTTCTTTTCTCT - 3'). F1 rollers were screened by PCR using the following primers (5' –CGCCAAATAAAGGACATTATGTAGCT TAT - 3' and 5'- TGACAAGTAAGTAAGCTGTACCTAATTTTC - 3'). Successful insertion resulted in a larger fragment and was again verified by sequencing.

An NLS was added to the DAM1284 TurboID strain using CRISPR-Cas9 gene editing as previously described, using Cas9 protein (0.25µg/µL), tracrRNA (0.02µg/µL, and crRNA (0.02µg/µL) (5'- GACGTCGTATGGGTAGCCAC [NGG]-3'), dpy-10(cn64) (40ng/µL). A ssODN (0.11µg/µL) was used as the repair template (5' – CCCAGGTCACCGTCTCCAGCAGCTCCA CCGGTGGCCCAGCCGCCAAGCGTGTCAAGCTCGACTACCCATACGACGTCCCAGACTAC GCCTACCCATA - 3'). F1 rollers were screened by PCR using the following primers (5'- GGGT GATCGTAGCTCCTATGAAG-3' and 5'- AAG GGCGATAAGCTTAAGTGGG - 3'). Successful insertion resulted in a larger fragment and was verified by sequencing.

**Immunofluorescence Methods**

The following primary antibodies were used in this study: chicken anti-HTP-3 (1:500, ref. 3), mouse anti-GFP (1:500, Roche), rabbit anti-GFP (1:500, ref. 4), rabbit anti-MSH-5 (1:10,000, SDIX), rat anti-RAD-51 (1:200, ref. 5), guinea pig anti-SUN-1 pS8 (1:1000, ref. 6), guinea pig anti-SYP-1 (1:200, ref. 7)), guinea pig anti-ZHP-3 (1:500; ref. 8), rabbit anti-REC-8 (1:1,000; Novus Biologicals), mouse anti-FLAG (1:200, Sigma), rat HIM-8 (1:500; ref. 9), mouse anti-αTubulin-FITC (1:500, DM1α, F2168; Sigma-Aldrich), rabbit anti-ASPM-1 (1:3000; ref. 10), mouse anti-HA.11 (1:100, clone 16B12, Biolegend), and rat anti-HA (1:2000, clone 3f10, Sigma Aldrich).

Secondary antibodies used were Alexa Fluor 488-, 555-, and 647-conjugated goat antibodies raised against the appropriate species (all used 1:200, Life Technologies).

Fixation for immunofluorescence used methods adapted from (11). 20-40 adults (24-28 hours post-late L4 stage) were dissected in egg buffer and fixed on charged slides (Fisher Scientific, Cat# 1255015) in 1% PFA in egg buffer (11) with 0.1% tween for 5 min at RT, immersed in liquid $N_2$ until frozen, and transferred to -20 ºC methanol for 5 min. For immunostaining of whole mount gonads, slides were then processed as in (11).

Nuclear spreads were prepared as in (12, 13), using a 10% v/v Hanks Balanced Salt Solution (HBSS, Life Technologies, 24020-117) for the dissection buffer.

For immunolocalization of PARG-1::GFP, we used a modified fixation procedure as described in (14) in which worms were fixed in 2.5% PFA for 2 mins at RT and then freeze-cracked in liquid nitrogen. Slides were placed in absolute ethanol at −20 °C for 10 mins and then washed in 1× PBST before proceeding as in (11).

For visualization of meiotic spindles, embryos released from gravid hermaphrodites were fixed and stained as in (15). To induce metaphase-arrested spindles (Fig S4B), worms were subjected to *emb-30 RNAi* as in (15).

#### FISH experiments and quantification

For FISH experiments, fixation methods were as described above. We used a set of oligopaint probes targeting a 1 Mb segment of chromosome II (genomic coordinates 11,500,001-12,500,001) (12). Slides were processed as in (16), using methods adapted from (17). Slides were mounted with vectashield (Vector Laboratories, Cat#H-1000) using No. 1.5 cover slips (Fisher Scientific, Cat# 12541019). Images and quantification used only the labelled fiducial probe.

Images of stitched whole-mount gonads were cropped and rotated so that meiotic progression proceeded from left to right along an x axis. Z-stack images comprising a single layer of non-overlapping germ cell nuclei were manually segmented using DAPI signals and polygon-shaped regions of interest (ROIs) were saved using FIJI software. Nuclei that partially overlapped other nuclei or ambiguous nuclei were not included in ROIs. An overlay of collected ROIs was used to clear the area outside of ROIs of signal throughout the stack. We then used the 3D Maxima Finder plugin in FIJI to identify FISH foci, generating output files of peak brightness and 3D positions of maxima (18). 3D Maxima Finder parameters used were 'Radius xy' = 3, 'Radius z' = 3, and 'Noise' = 100. The 'Minimum peak Height' parameter was determined empirically, using the maximum value of background fluorescence and iteratively running the plugin to minimize false positive and false negative foci identified.

From the output file of 3D Maxima Finder, foci were assigned to individual nucleus ROIs based on position, using a custom Python script (19). Assignments were checked manually to remove overlap or nuclei without any foci before final foci counts were plotted using GraphPad Prism. The xy position of each ROI was used to approximate its location in four 'zones' of the germ line. Germline zone segmentation was done using only the DAPI channel. Zone 1 was defined to span from the distal end of the germ line until the point where germ line width increased around transition zone entry. Zone 4 was defined as beginning 10 rows of nuclei prior to the point where the germline decreased to a width of one to two nuclei per cell row. Zone 2 was defined as the first one-third of the distance between the end of zone 1 and the start of zone 4, and Zone 3 was defined as nuclei within the remaining two-thirds of that distance (Figure S3).

#### Imaging

All images of whole mount gonads and nuclear spread preparations were acquired on a DeltaVision OMX Blaze microscope with a 100 x 1.4 NA widefield objective and z-stacks spaced at 200nm. Images were deconvolved and registration corrected with SoftWoRx and, where appropriate, stitched together using the 'Grid/Collection stitching' FIJI plugin (20). Displayed images are maximum intensity projections of z-stacks encompassing whole nuclei, except where noted.

Images of PARG-1::GFP in the germ lines of live whole worms were acquired using a Zeiss Axioimager microscope with a 40 x air 0.9 NA widefield objective.

Images of meiotic spindles in Figure S4 were acquired at 4°C on Zeiss LSM 880 Laser Scanning Confocal Microscope with a 63x 1.58 NA objective; Z-stacks were obtained at 300nm increments. Images shown are maximum intensity projections processed in FIJI. Polar bodies numbers in post-meiotic embryos were assessed using images acquired with a Zeiss Axioimager microscope with a 40 x 0.9 air NA widefield objective.

**TurboID sample preparation**

*Growth of E. coli OP50 and preparation of OP50-Seeded NGM Plates:*

To prepare OP50 for nematode culture, 5 mL of an overnight OP50 culture was inoculated into 1 L of LB medium in wide-bottom 2 L flasks and incubated at 37°C with shaking overnight. Following incubation, cultures were pelleted using a floor centrifuge at 6000 rpm for 10 min at room temperature. The supernatant was discarded, and pellet was resuspended in 100 mL of water and frozen at -20°C. For seeding, frozen OP50 stocks were thawed in a microwave (10–30 s intervals, with intermittent shaking to disperse clumps) and 2–3 mL of thawed OP50 suspension was applied per 15 cm NGM agar plate to ensure dense and uniform bacterial lawns.

*Preparation and harvesting of worm cultures:*

Synchronized L1 worms (21) were plated at 20-25,000 worms per 15 cm plate. Worms were incubated at 20°C for 72 hours, gravid worms were harvested into 15 mL conical tubes, pelleted by centrifugation, and washed three times in M9 buffer (22).

Samples were nutated for 10 min in M9 to allow clearance of gut contents, then washed once more with Milli-Q water to remove residual salts. After final centrifugation, the supernatant was carefully aspirated, and worm pellets were snap-frozen in liquid nitrogen and stored at −80°C.

*Cryogenic Grinding and lysis:*

Frozen worm pellets (~400–500 µL) were ground using a Biospec Cryo-Cup Grinder (Cat#206). Each pellet was transferred into a liquid nitrogen-precooled cryo-cup, and ground to a fine powder with a chilled pestle. The powder was transferred to pre-chilled 50 mL conical tubes and stored at −80°C, or processed immediately.

Worm powder was resuspended in 1.5 mL of chilled nuclear lysis buffer (50 mM Tris-HCl pH 8.0, 5 mM $MgCl_2$, 1 mM EGTA, 150 mM NaCl, 10% glycerol). Samples were vortexed intermittently (for 20-30 sec intervals, followed by 1 min rest on ice) until homogenized, then incubated on ice for 5 min. Benzonase (1.5 µL per sample) was added, and samples were nutated for 5 min at 4°C. Crude lysate (30 µL) was saved, and samples were centrifuged at 2000 × g for 10 min. A second 30 µL aliquot of the clarified supernatant was saved as input. The supernatant was transferred to new tubes, avoiding the fatty layer.

*Protein Quantification and Affinity Purification:*

Protein concentrations were determined using a Bradford assay. Sample volumes were adjusted to yield 1200 µg total protein per sample.

For each sample, 150 µL of streptavidin magnetic beads (Thermo Scientific, Ref# 88817) were equilibrated in 500 µL of lysis buffer using a magnetic rack. After removal of the equilibration buffer, clarified protein lysates were added to the beads and nutated with beads overnight.

*Sequential washing of magnetic beads:*

All wash steps were performed at 4°C to preserve protein integrity. Samples were subjected to a sequential wash protocol with ice-cold solutions. At each step, 1 mL of the indicated solution was added to the sample, mixed gently, and magnetized. The supernatant was discarded, and the beads were retained for the next wash. The washing sequence was as follows: 1 M KCl, 0.1 M $Na_2CO_3$, 2 M urea, 4 M urea, nuclear lysis buffer (x2), and seven consecutive washes with PBS.

**Mass Spectrometry sample preparation and analysis for TurboID**

*Sample Preparation:*

Nine biotinylated protein samples (three replicates per condition) were digested on streptavidin-coated magnetic beads. The proteins were reduced with dithriothreitol (DTT) added to a final concentration of 10mM and incubated for five minutes at 55°C, followed by head-over-head mixing at room temperature for 25 minutes using a Barnstead Thermolyne Labquake rotisserie shaker. They were then alkylated with acrylamide added to a final concentration of 30 mM, at room temperature for 30 minutes. This was followed by overnight digestion at 37°C using 500 ng of mass spectrometry grade trypsin/LysC mix (Promega, Madison, WI). Post-digestion, samples were quenched with formic acid (adjusted to a pH ~3) and desalted using MonoSpin C18 Solid-Phase Extraction (SPE) columns (GL Sciences, Torrance, CA). Finally, the samples were dried via SpeedVac (ThermoFisher Scientific, San Jose, CA) and exchanged into LC-MS reconstitution buffer (2% acetonitrile with 0.1% formic acid in water) for instrumental analysis.

*LC-MS/MS Analysis:*

Proteolytically digested peptides were separated using an in-house pulled and packed reversed phase analytical column (~25 cm in length, 100 microns of I.D.), with Dr. Maisch 1.9-micron C18 beads as the stationary phase. Separation was performed with an 80-minute reverse-phase gradient (2-45% B, followed by a high-B wash) on an Acquity M-Class UPLC system (Waters Corporation, Milford, MA) at a flow rate of 300 nL/min. Mobile Phase A was 0.2% formic acid in water, while Mobile Phase B was 0.2% formic acid in acetonitrile. Peptide cations were formed via electrospray ionization and analyzed by an Orbitrap Exploris 480 mass spectrometer (ThermoFisher Scientific). The mass spectrometer was operated in a data-dependent mode using HCD fragmentation for MS/MS spectra generation.

The .RAW data were analyzed using Byonic v5.2.5 (Protein Metrics, Cupertino, CA) to identify peptides and infer proteins. A concatenated FASTA file containing Uniprot *Caenorhabditis elegans* proteins, bait sequences, and other likely contaminants was used to generate an *in silico* peptide library. Proteolysis with Trypsin/LysC was assumed to be semi-specific allowing for N-ragged cleavage with up to two missed cleavage sites. The precursor and fragment ion tolerances were set to 12 ppm. Cysteine capped with propionamide was set as a fixed modification in the search. Variable modifications included oxidation on methionine, histidine, and tryptophan, deamidation of glutamine and asparagine, glutamine and glutamic acid cyclization, and lysine biotinylation. Proteins were held to a false discovery rate of 1% using the standard reverse-decoy technique (23).

**PARG-1::GFP Immunoprecipitation and Mass Spectrometry:**

*Sample preparation for mass spectrometry analysis:*

Fractionation and immunoprecipitation of PARG-1::GFP were performed as in (24), using Chromotek gta-20 GFP-TRAP beads. For each immunoprecipitate, both on-bead digestion and elution protocols for preparing peptides were followed (below). As PARG-1 peptides were only detected in the on-bead digestion samples, only the data from the on-bead samples were considered.

For on-bead digestion, the beads were resuspended in 30 µL 2 M urea and 50 mM ammonium bicarbonate. Disulfide bonds were reduced with 1.5 µL of 200 mM dithiothreitol (DTT) for 30 min at room temperature before adding 1.5 µL of 500 mM iodoacetamide and incubating for 15 min at room temperature in the dark. The remaining iodoacetamide was quenched with 0.75 µL of 200 mM DTT for 10 min. Proteins were digested with 150 ng trypsin (Trypsin Gold, Promega) in 1.5 µL 50 mM ammonium bicarbonate at room temperature for 90 minutes. The supernatant was transferred to a new tube. The beads were rinsed with 30 µL 2 M urea and 50 mM ammonium bicarbonate and pooled with the previous supernatant. The combined supernatants were diluted with 50 mM ammonium bicarbonate to reach a urea concentration of 1 M. Then the solution was digested further with another 150 ng trypsin (Trypsin Gold, Promega) in 1.5 µL 50 mM ammonium bicarbonate at 37°C overnight. The digest was stopped by the addition of 10% trifluoroacetic acid (TFA) to a final concentration of 0.5%, and the peptides were desalted using C18 Stagetips (25).

For elution, the beads were washed once with and 50 mM ammonium bicarbonate, the supernatant removed and then resuspended in 20 µL 0.1% TFA and incubated for 3 min. The supernatant was transferred to a new tube and the elution step was repeated another three times. After adjusting the pH with 1 M Tris pH 8.5, 8 M urea solution was added to reach a final concentration of 2 M. Reduction and alkylation with DTT and IAA was performed as above, and the proteins digested with 200 ng trypsin at 37°C overnight. The digest was stopped by the addition of 10% trifluoroacetic acid (TFA) to a final concentration of 0.5%, and the peptides were desalted using C18 Stagetips (25).

*Liquid chromatography-mass spectrometry analysis:*

LC-MS analysis was performed on an UltiMate 3000 RSLCnano LC system (Thermo Scientific) coupled to a Q Exactive HF-X hybrid quadrupole-Orbitrap mass spectrometer (Thermo Scientific). The system was equipped with a nano-spray Flex ion-source (Thermo Scientific) using coated emitter tips (New Objective).

Peptides were loaded onto a trap column (Acclaim PepMap C18 5 µm 300 µm x 5mm, Thermo Scientific) using 0.1% TFA as mobile phase, and separated on an analytical column (Acclaim PepMap C18, 50 cm × 0.75 mm, 2 µm, Thermo Scientific), applying a linear gradient starting with a mobile phase of 98% solvent A (0.1% FA) and 2% solvent B (80% acetonitrile, 0.08% FA), increasing to 35% solvent B over 120 min at a flow rate of 230 nl/min. The analytical column was heated to 30°C.

The mass spectrometer was operated in data-dependent mode, survey scans were obtained in a mass range of 380-1650 m/z with lock mass activated, at a resolution of 120k at 200 m/z and an AGC target value of 3E6. The 10 most intense ions were selected with an isolation width of 2 Da, fragmented in the HCD cell at 27% collision energy and the spectra recorded at a target value of 1E5 and a resolution of 30k. Peptides with a charge of +1 or >+6 were excluded from fragmentation, the peptide match feature was set to preferred, the exclude isotope feature was enabled, and selected precursors were dynamically excluded from repeated sampling for 30 seconds.

*Data analysis:*

Raw data were processed using the MaxQuant software package (version 1.5.5.1, ref. 26)) and the Uniprot *C. elegans* reference proteome (downloaded 2017-01-09) as well as a database of most common contaminants (MaxQuant contaminant file). The search was performed with full trypsin specificity and a maximum of two missed cleavages at a protein and peptide spectrum match false discovery rate of 1%. Carbamidomethylation of cysteine residues were set as fixed, oxidation of methionine, and N-terminal acetylation as variable modifications. For label-free quantification the “match between runs” feature and the LFQ function were activated (27) - all other parameters were left at default.

MaxQuant search results were further processed using the Perseus software package (version 1.5.5.3, ref. 28). Contaminants, reverse hits, and proteins identified only by site were removed and the log2 transformed LFQ values were used for protein quantification. Data were median normalized, missing values were replaced with a fixed value close to the detection limit, and log2 ratios were calculated for bait / control.

#### Supplemental Figure Legends

**Figure S1. *duo-1* males have defects in spermatogenesis.** Top: Images of *WT* whole-mount male gonad immunostained for RAD-51, HTP-3, SYP-1, and co-stained with DAPI. Insets show early pachytene nuclei (yellow box), mid/late pachytene nuclei (red box), and mature spermatid nuclei, which have a uniform, rounded appearance (white box). Bottom: *duo-1* mutant gonad stained as above, showing hyperaccumulation of RAD-51 and defective SC assembly (red and yellow insets). Large inset of DAPI-stained nuclei shows non-uniform and aberrantly shaped nuclei in the proximal germ line, where mature spermatids are found in *WT*. Arrows indicate examples of aberrant chromatin in the mature spermatid region, including elongated DNA structures (open arrows) and small spots that may correspond to chromosome fragments and/or incomplete chromosome complements (closed arrows). Scale bars = 10μm.

**Figure S2. Chromatin morphology and assembly/stability of other SC components and associated proteins in *duo-1* mutants.** *A)* Images of DAPI-stained chromatin in the *WT* and *duo-1* mutant gonads depicted in Fig. 1, showing nuclei exhibiting clustered chromatin organization in the *duo-1* mutant as well as a compact subset of chromatin that persists into the mid-pachytene region. Insets show (from left to right) nuclei in the early, mid- and late pachytene regions. Scale bars = 10μm. *B)* Immunostaining for axis protein HIM-3 and SC-associated RING finger protein ZHP-3, showing a *duo-1* phenotype similar to that shown in Figure 1. Scale bars = 10μm.

**Figure S3. X chromosome pairing in *duo-1* mutants.** Immunostaining for X-chromosome pairing-center protein HIM-8, showing that X chromosomes pair in the *duo-1* mutant and that the X chromosome remains tightly compacted through the mid/late pachytene region. Right insets showing HIM-8 in cyan and DAPI in red highlight compact X chromosome territories. Scale bars = 10μm.

**Figure S4. Failure of second polar body extrusion and abnormal oocyte spindle and chromosome structures in *duo-1* mutant embryos.** *A)* Left: Sample images of polar bodies (asterisks) detected by DAPI staining in early embryos. Right: Graph depicting quantitation of polar bodies in 1-2 cell post-meiotic embryos, showing that two polar bodies were detected in most wild-type embryos (n = 50), whereas only a single polar body was detected in most duo-*1* mutant embryos (n = 50). The single polar body phenotype is a hallmark of mutants that exhibit premature loss of REC-8 cohesin (29), where it is indicative of (roughly) equational segregation of sister chromatids occurring at meiosis I coupled with failure of polar body extrusion at meiosis II. Failure to extrude a second polar body after equational segregation at meiosis I will result in a (near) diploid chromosome complement in the female pronucleus, increasing the likelihood of a viable, near euploid chromosome complement in the zygote. This can explain the relatively high incidence of embryonic hatching in *duo-1* mutants relative to meiotic mutants that lack chiasmata and undergo homolog mis-segregation at meiosis I. *B, C)* High-resolution confocal images of oocyte meiotic spindles in metaphase I-arrested *B)* and unarrested *C)* embryos immunostained for $\alpha$-tubulin and spindle-pole marker ASPM-1 and counterstained with DAPI to visualize chromosomes. Example metaphase I-arrested spindles from the *duo-1* mutant shown in *B)* are less symmetric than control metaphase I-arrested spindles, and chromosomes are more elongated and less compact. For unarrested embryos/spindles in C): while it was straightforward to capture meiotic spindles representing standard meiotic division stages (metaphase I, anaphase I, metaphase II and anaphase II) in wild-type embryos, it was difficult to locate and/or unambiguously assign spindle stages in *duo-1* mutant embryos. Moreover, we frequently detected unusual structures that we have never observe in wild-type controls, *e.g.* several instances of an apparent maternal pronucleus attached *via* a tubulin-enriched structure (presumably a persistent spindle midbody) to an apparent disk-shaped polar body flattened up against the cortex (far right bottom panel in C).

**Figure S5. RAD-51 accumulation in the *duo-1* mutant is SPO-11 dependent, while diakinesis decompaction phenotype is not.** *A)* Representative images of proximal oocytes from *WT*, *spo-11*, *duo-1* and *spo-11; duo-1* backgrounds, showing that the *duo-1* mutant decompaction phenotype is not SPO-11 dependent. Scale bar = 5μm. *B)* Immunostaining of RAD-51, showing that RAD-51 accumulation in a *duo-1* background is SPO-11 dependent. Scale bar = 10μm. *C)* Immunostaining for RAD-51 and SYP-1, showing that polycomplexes are present in *spo-11; duo-1* mutants. Scale bar = 10μm.

**Figure S6. Immunostaining in nuclear spread preparation.** *A)* Reduced HTP-3 intensity in *duo-1* spread nuclei. Greyscale images from Fig. 3C, showing intensity-matched HTP-3 staining optimized for WT (left) or for *duo-1* (right). *B)* Images of nuclear spreads co-stained for MSH-5 and COSA-1. Scale bars in all panels = 10μm.

**Figure S7. Extension of CHK-2 active zone in *duo-1* mutants***.* *A)* Immunostaining of whole mount *WT* and *duo-1* germ lines for SUN-1pS8, a marker of CHK-2 activity in *C. elegans* meiosis. Scale bar = 10μm. *B)* Quantification of length of CHK-2 active zone in *WT* and *duo-1*, showing a significant increase in length in *duo-1* mutant germ lines. For quantification, germ lines were straightened and measured for: i) length from meiotic onset to end of the pachytene region, and ii) length of the meiotic SUN-1p8-stained region, starting from meiotic onset to the last row in which the majority of nuclei showed bright SUN1-p8 staining. 3 gonads were scored per genotype; error bar indicates SEM, *** indicates p = 0.0007, two-tailed unpaired T-test.

**Figure S8. Extended depletion of DUO-1 in whole gonads***.* Immunostaining of whole mount *WT* and *aid::duo-1::aid* germ lines shown in Figure 4, showing breakdown of SC into aggregates by diplotene, similar to the *duo-1* mutant phenotype. Scale bar = 10μm.

**Figure S9. Identification of PARG-1 as an interactor of DUO-1.** *A-C)* One-sided volcano plots of DUO-1::GFP and NLS::TurboID compared to A) both controls, B) DUO-1::GFP only control, and C) NLS::TurboID only control. Each point represents a protein identified by LC-MS from biotinylated proteins captured by streptavidin pull down and purification. x-axis shows $log_2$ fold difference in abundance between indicated conditions and y-axis shows -log 10 of the p value. Stringent significance curves were generated in Perseus (FDR = 0.05, s0 = 0.1). Blue points correspond to meiotic HORMAD proteins and pink points correspond to cohesin components. See Methods for full details and strain genotypes. *D)* Nuclear localization of GFPnanobofy:NLS::HA:TurboID with and without DUO-1:GFP in control and experimental strains used for TurboID experiments, showing nuclear localization of HA. *E)* PARG-1::GFP fluorescence in live worms. *F)* List of top hits from co-immunoprecipitation/MS experiment targeting PARG-1:GFP. Inclusion criteria: $\geq$ 5 peptides in experimental IP, experimental/control ratio $\geq$ 3.

#### Figure S1

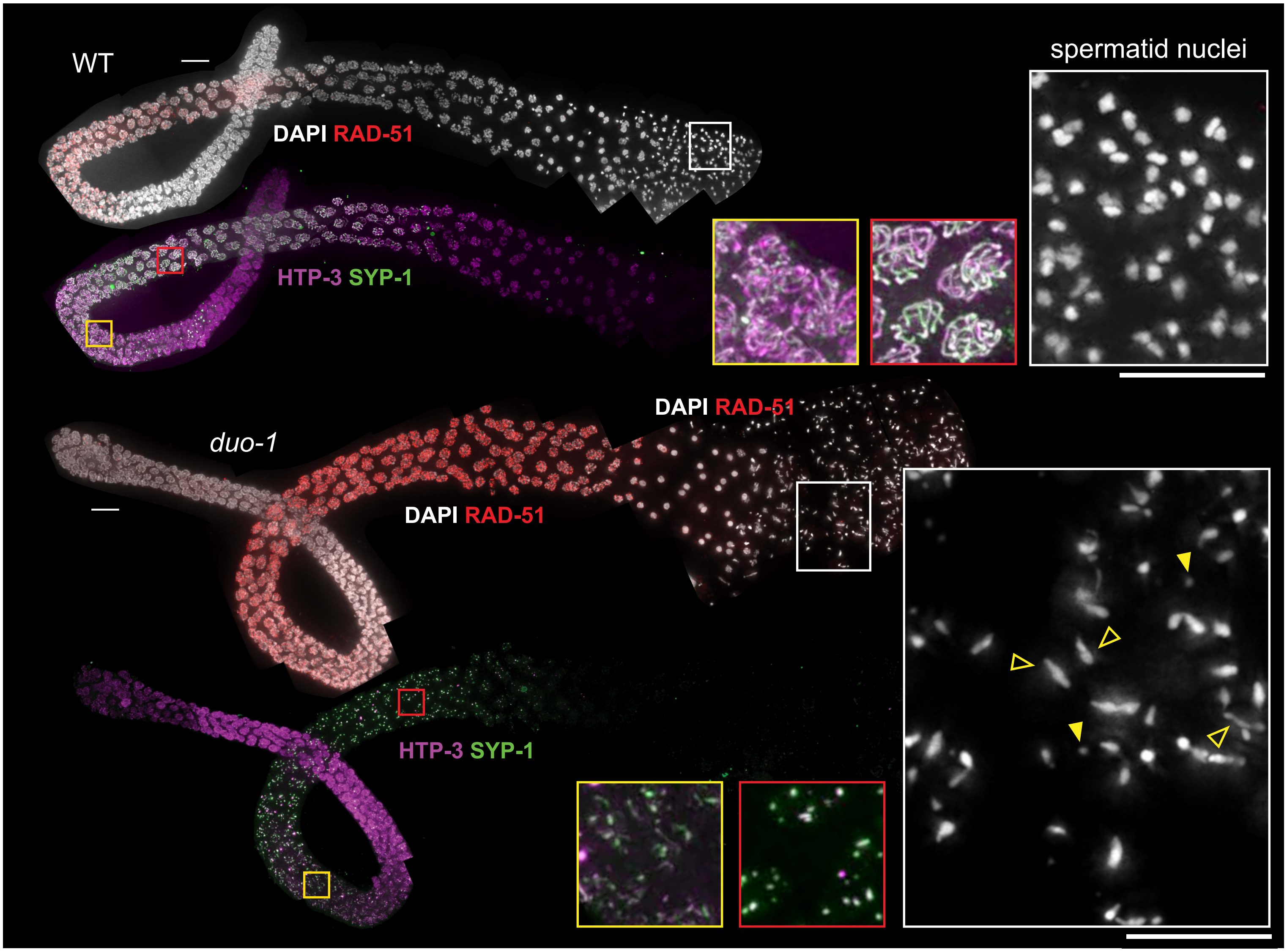
WT
DAPI RAD-51
HTP-3 SYP-1
spermatid nuclei
duo-1
DAPI RAD-51
DAPI RAD-51
HTP-3 SYP-1

### Figure S2

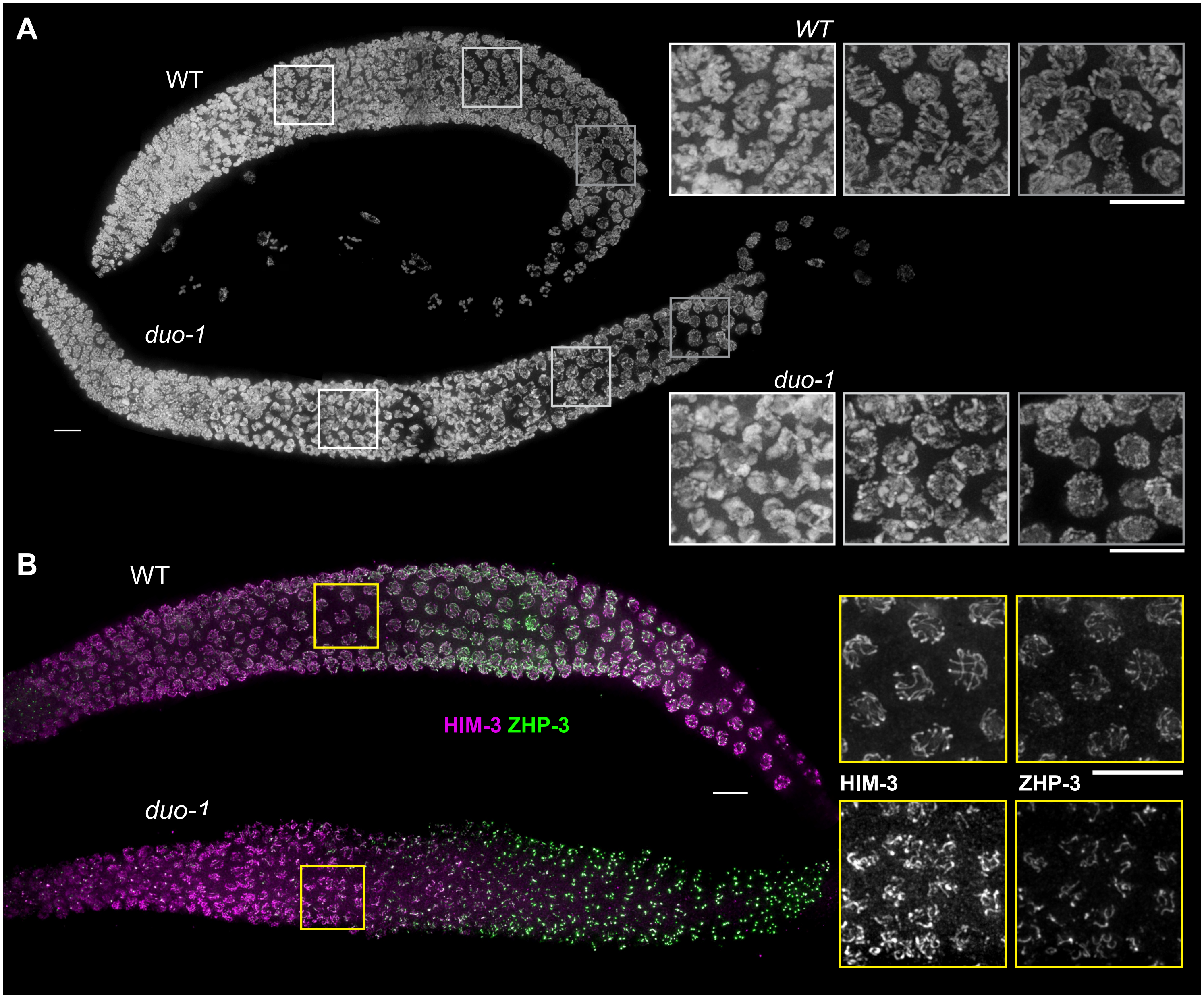

#### Figure S3

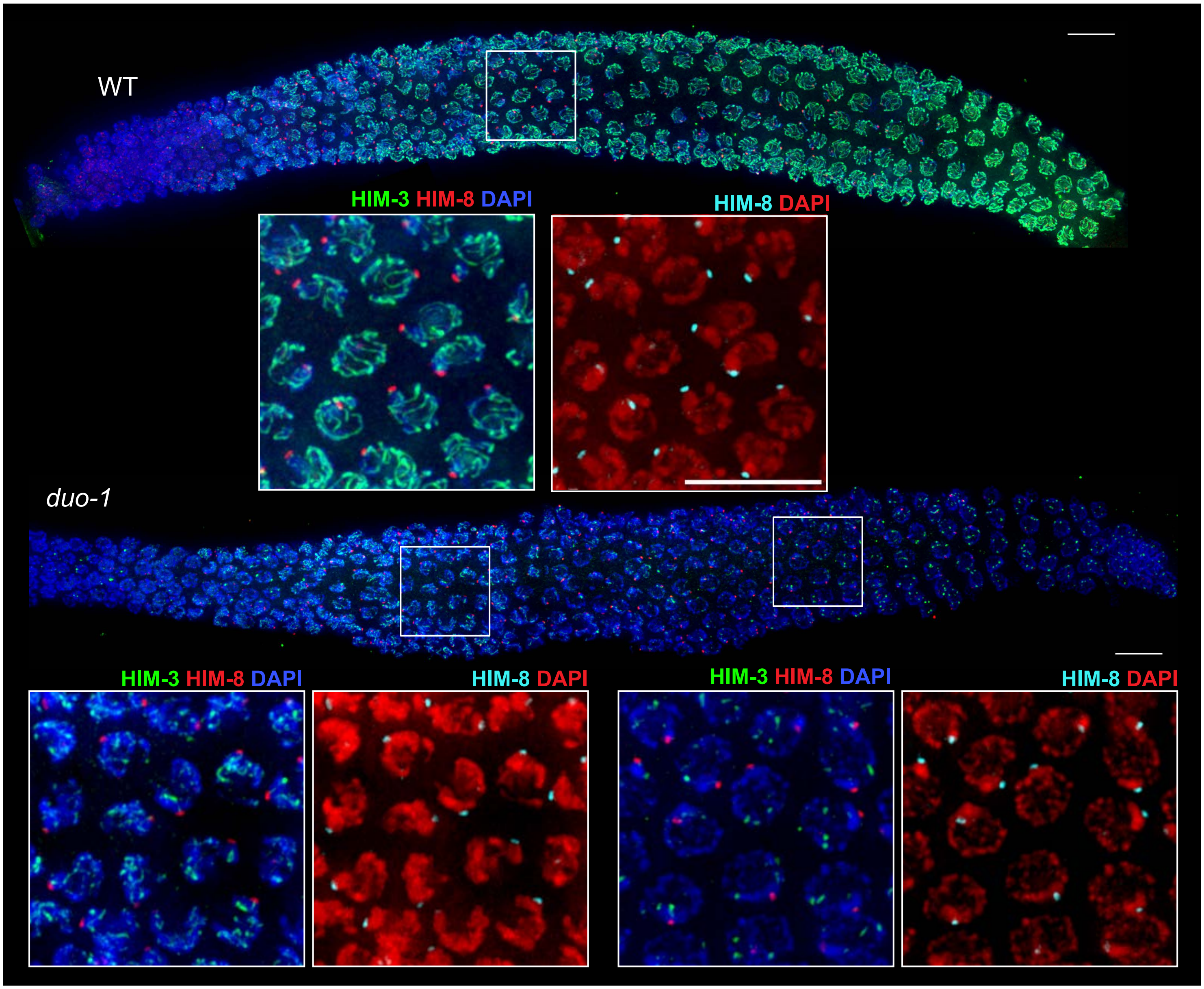

WT
HIM-3 HIM-8 DAPI
HIM-8 DAPI
duo-1
HIM-3 HIM-8 DAPI
HIM-8 DAPI
HIM-3 HIM-8 DAPI
HIM-8 DAPI

#### Figure S4

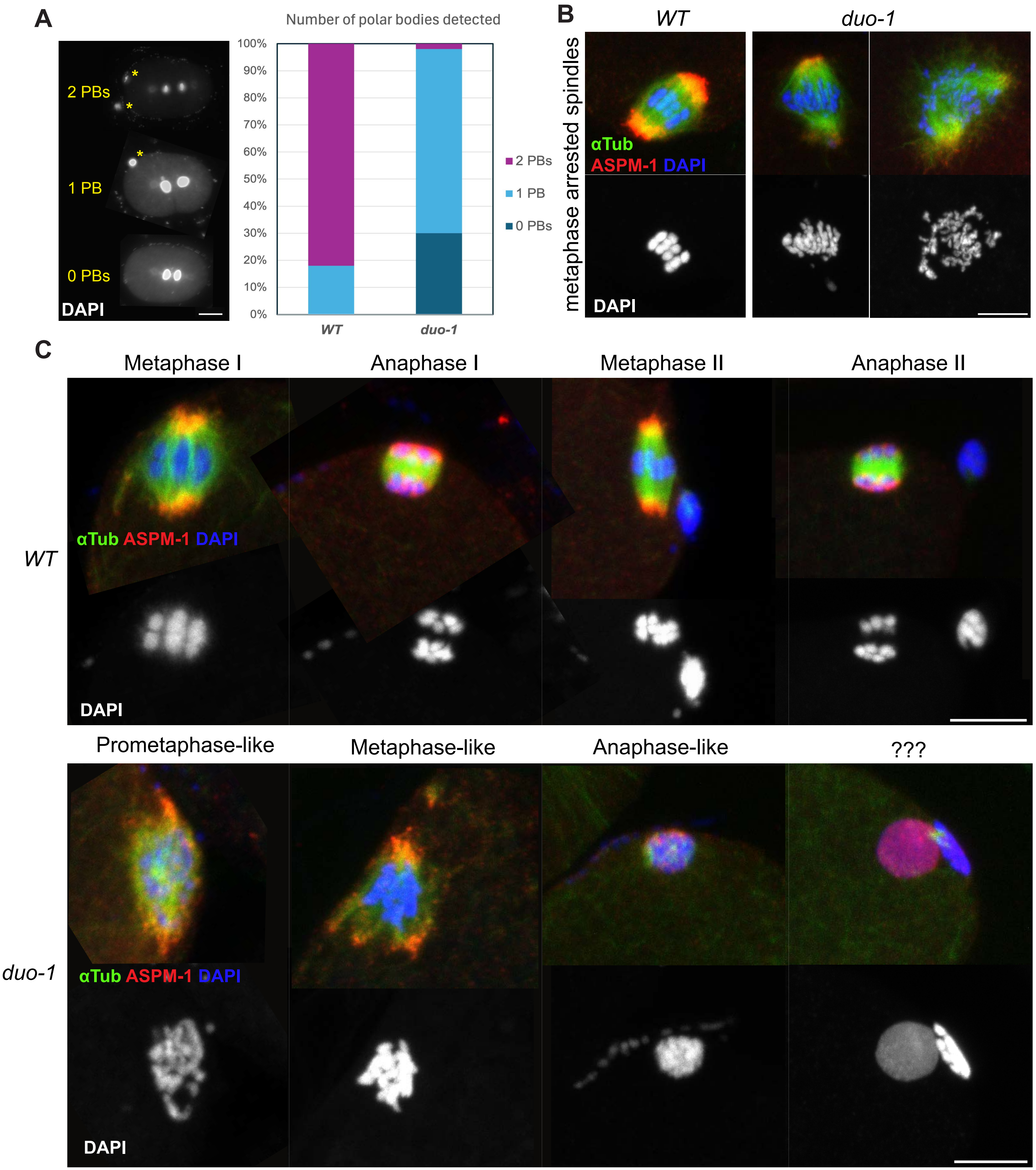
A
2 PBs
1 PB
0 PBs
DAPI
Number of polar bodies detected
100%
90%
80%
70%
60%
50%
40%
30%
20%
10%
0%
WT
duo-1
2 PBs
1 PB
0 PBs
B
WT
duo-1
metaphase arrested spindles
αTub
ASPM-1 DAPI
DAPI
C
Metaphase I
Anaphase I
Metaphase II
Anaphase II
WT
αTub ASPM-1 DAPI
DAPI
Prometaphase-like
Metaphase-like
Anaphase-like
???
duo-1
αTub ASPM-1 DAPI
DAPI

Figure S5

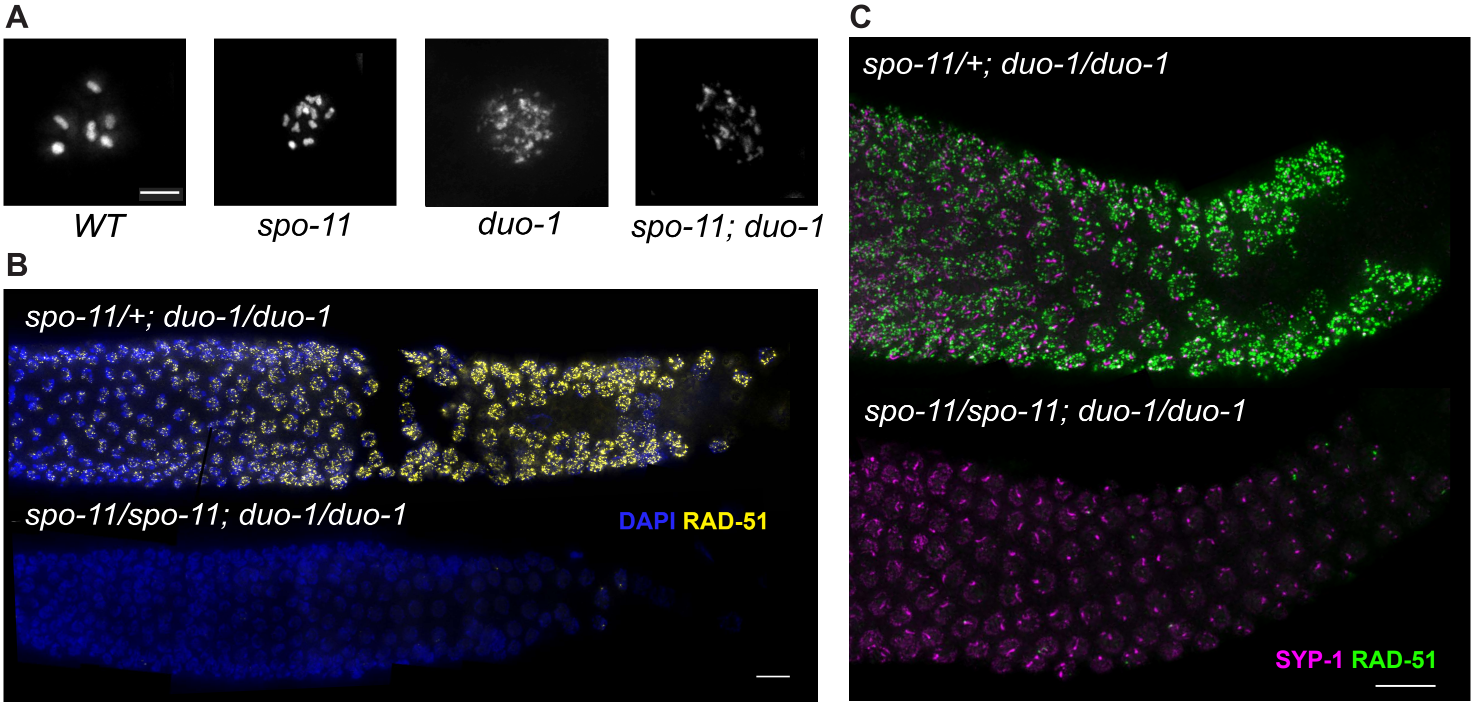
A
WT
spo-11
duo-1
spo-11; duo-1
B
spo-11/+; duo-1/duo-1
spo-11/spo-11; duo-1/duo-1
DAPI RAD-51
C
spo-11/+; duo-1/duo-1
spo-11/spo-11; duo-1/duo-1
SYP-1 RAD-51

#### Figure S6

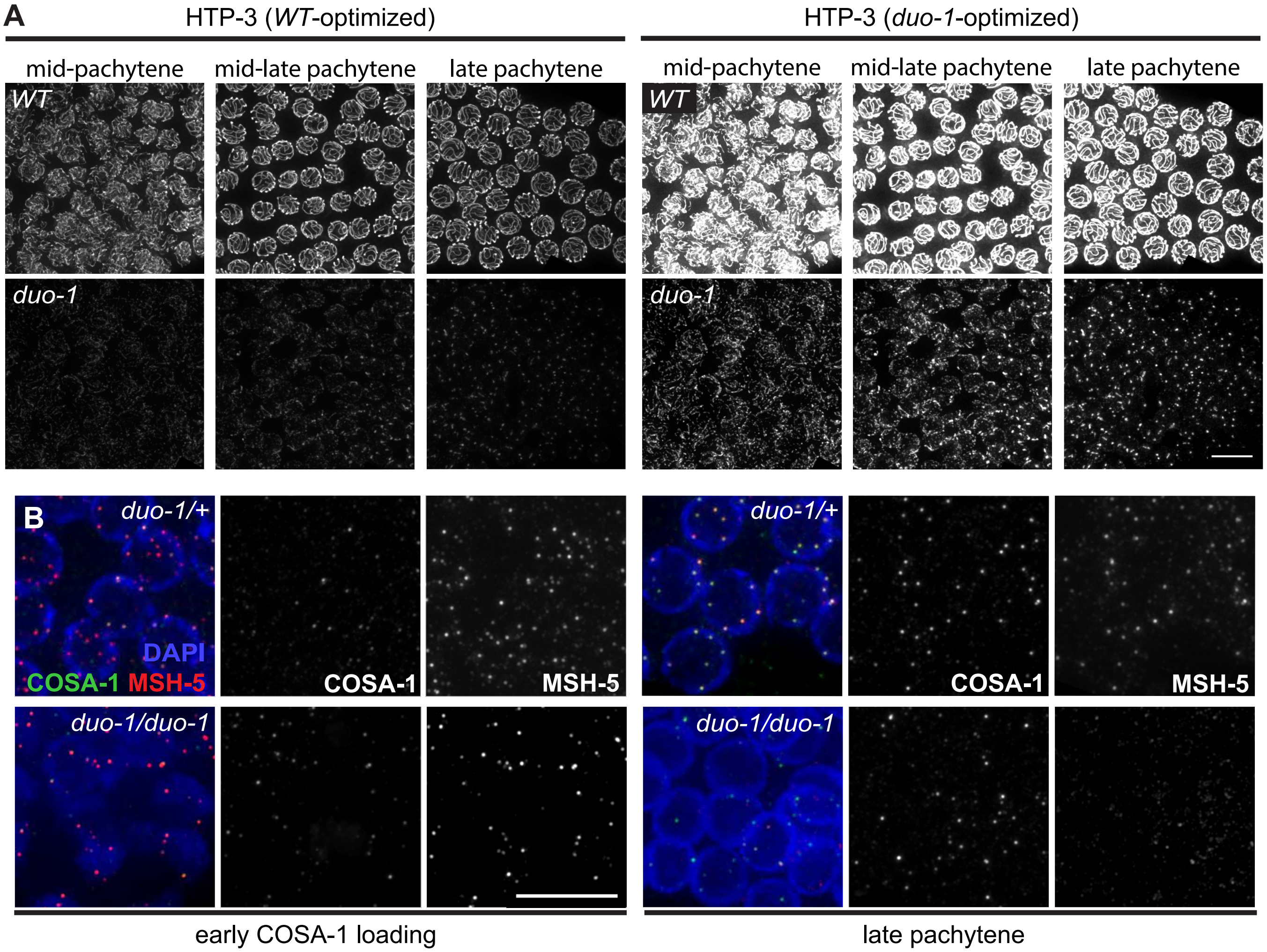
A
HTP-3 (WT-optimized)
HTP-3 (duo-1-optimized)
mid-pachytene
mid-late pachytene
late pachytene
WT
duo-1
B
duo-1/+
duo-1/duo-1
DAPI
COSA-1
MSH-5
early COSA-1 loading
late pachytene

#### Figure S7

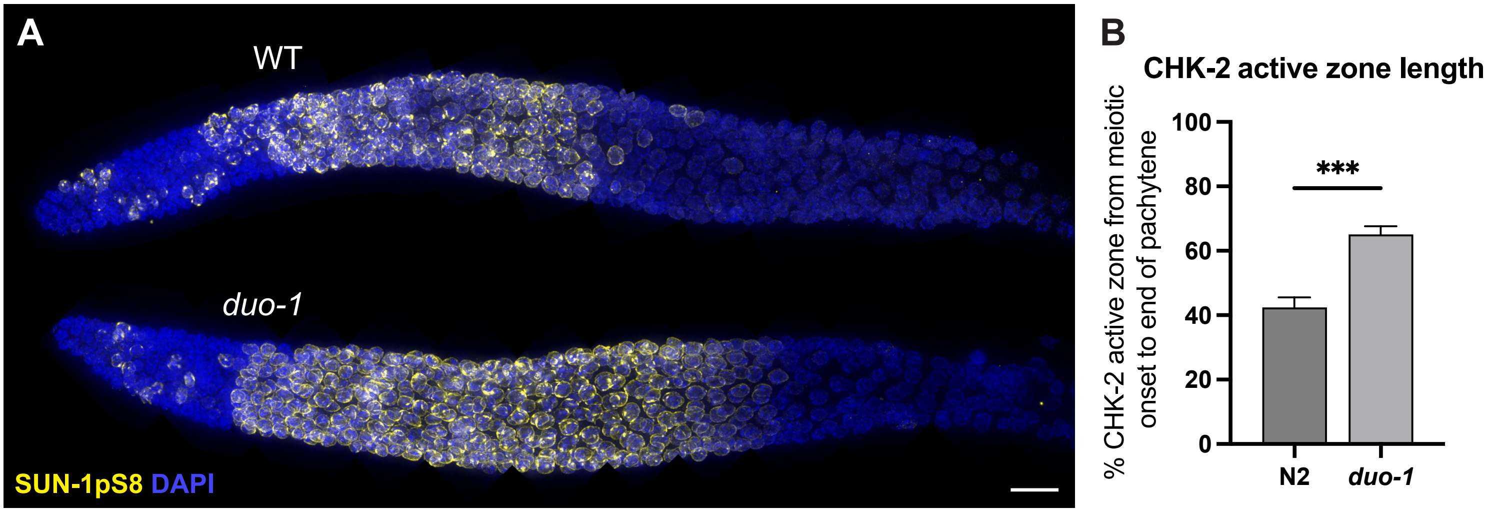

A
WT
duo-1
SUN-1pS8 DAPI
B
CHK-2 active zone length
% CHK-2 active zone from meiotic onset to end of pachytene
100
80
60
40
20
0
***
N2
duo-1

#### Figure S8

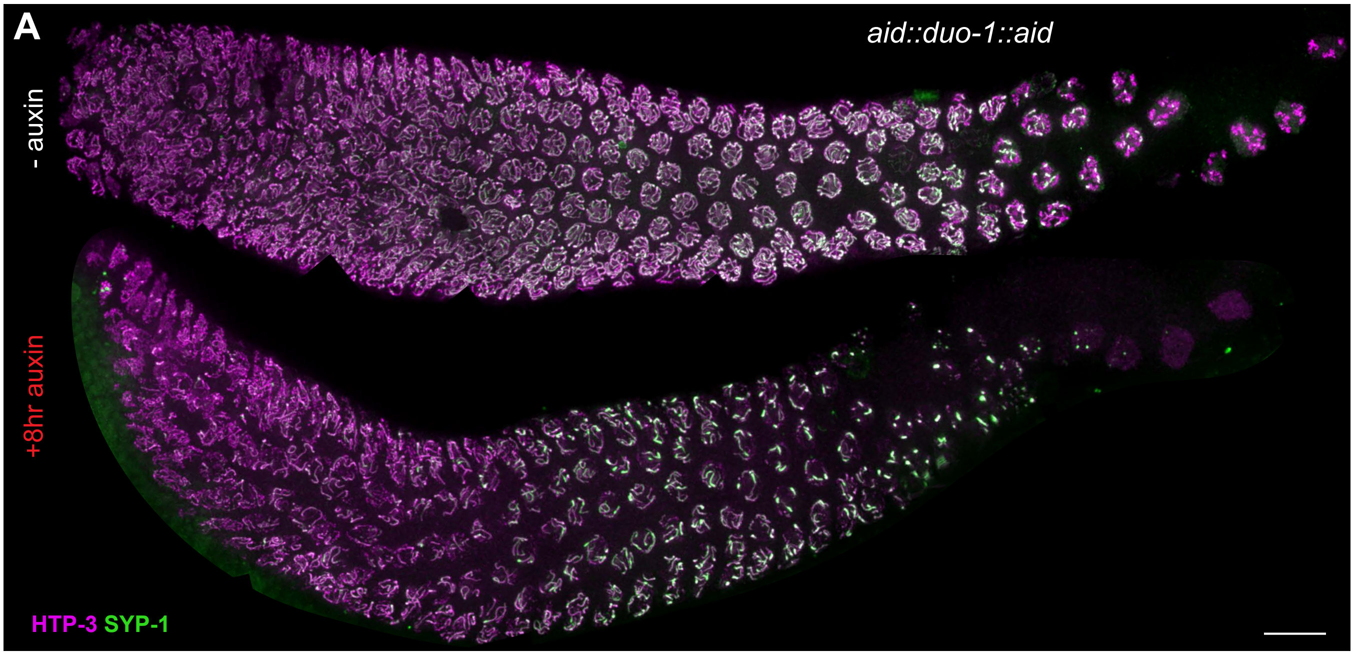

#### Figure S9

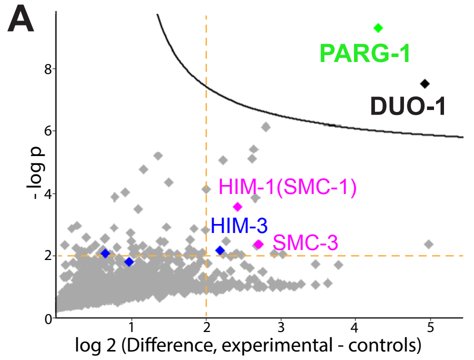


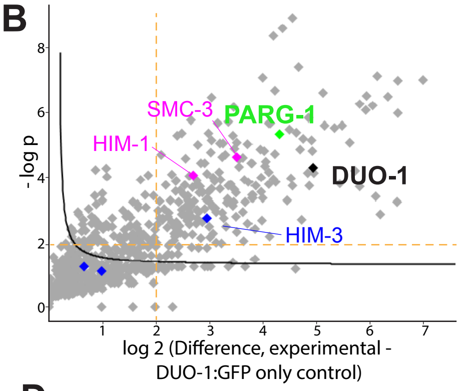


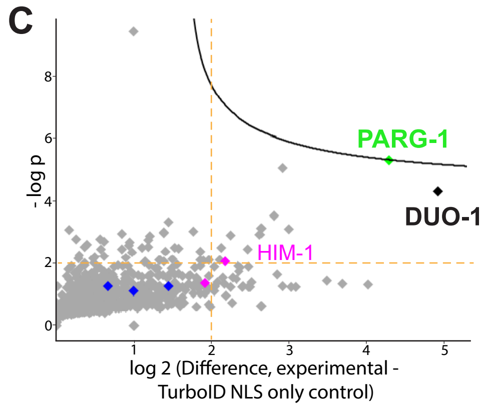


D

| Protein | Peptide counts from GFP IP/MS | |
|---|---|---|
| | control | *parg-1::GFP* |
| PARG-1 | 6 | 25 |
| HTP-1 | 1 | 10 |
| DUO-1 | 4 | 14 |
| COH-4 | 2 | 6 |
| SYP-1 | 5 | 15 |
| ADK-1 | 1 | 6 |

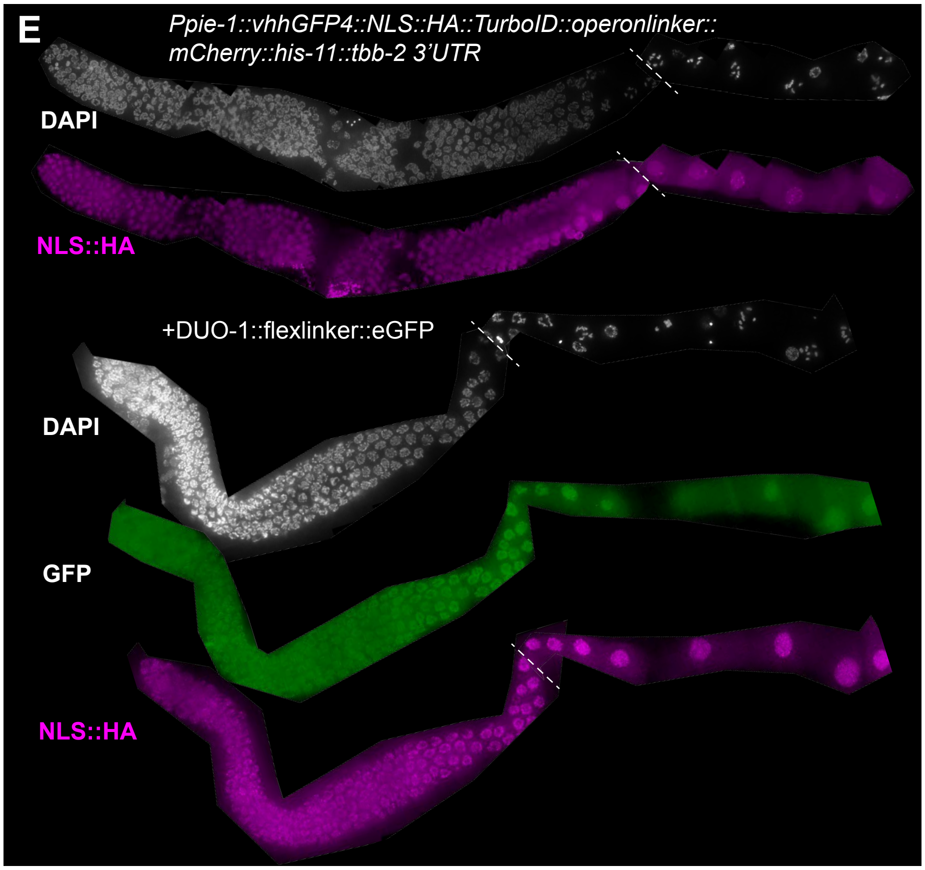
